# How tasks change whole-brain functional organization to reveal brain-phenotype relationships

**DOI:** 10.1101/870287

**Authors:** Abigail S. Greene, Siyuan Gao, Stephanie Noble, Dustin Scheinost, R. Todd Constable

## Abstract

Functional connectivity (FC) calculated from task fMRI data better reveals brain-phenotype relationships than rest-based FC, but how tasks have this effect is unknown. In over 700 individuals performing 7 tasks, we use psychophysiological interaction (PPI) and predictive modeling analyses to demonstrate that task-induced changes in FC successfully predict phenotype, and these changes are not simply driven by task activation. Activation, however, is useful for prediction only if the in-scanner task is related to the predicted phenotype. Given this evidence that tasks change patterns of FC independent of activation to amplify brain-phenotype relationships, we develop and apply an inter-subject PPI analysis to further characterize these predictive FC changes. We find that task-induced consistency of FC patterns across individuals is useful for prediction—to a point; these results suggest that tasks improve FC-based prediction performance by de-noising the BOLD signal, revealing meaningful individual differences in brain functional organization. Together, these findings demonstrate that, when it comes to the effects of in-scanner tasks on the brain, focal activation is only the tip of the iceberg, and they offer a framework to best leverage both task activation and FC to reveal the neural bases of complex human traits, symptoms, and behaviors.

## Introduction

Functional connectivity analyses have offered sweeping insights into the macroscale neural circuits underlying complex cognitive processes, finding these circuits to be broadly distributed across the human brain (e.g., ^1–5^). Such analyses are typically performed using resting-state data^6,7^, revealing “intrinsic connectivity networks” that recapitulate networks invoked during task execution^8^. This correspondence—along with demonstrations of the stability of functional connectivity (FC) patterns between resting and task states^9–11^—suggests that the functional network architecture of the human brain is relatively state-invariant. Nevertheless, there is a growing consensus that FC contains useful dynamic, rather than just static, information^12^. Task-induced changes in patterns of FC have been shown to be widely distributed across the brain^13^, to subserve the task at hand^14^, to make individuals more identifiable^15^, and to improve FC-based prediction of both task performance^1^ and stable traits, such as intelligence measures^16^. Together, these findings suggest that task-induced changes in FC, while perhaps low-amplitude and/or local perturbations of a core functional architecture, are functionally significant and may amplify individual differences in brain functional organization.

Thoughtfully leveraging such task-general changes therefore holds the promise of revealing the neural bases of a wide range of phenotypes, but the nature of these changes is not known. In particular, the question of whether task-induced FC changes reflect task-evoked activation, changes in neural interaction, or some combination of the two has received substantial attention. While some have raised concerns that inadequate removal of task-evoked activation from node time courses may yield spurious patterns of FC^17^, others have demonstrated that task-evoked activation and task-induced changes in FC can be cleanly dissociated^18^ and contain complementary information about task performance^19^, even when task-evoked activation is not removed from the BOLD signal^20^. Further, task-evoked FC (that is, task-induced changes in FC attributable to task-evoked activity) explains relatively little of the total task-induced change in FC^21^.

Here, we demonstrate that task-induced changes in FC reveal valuable, phenotype-relevant information, independent of any task-evoked activation, and use a range of analyses to characterize these changes. First, we sought to better understand why task FC-based models better predict phenotypic measures than rest FC-based models^16^: is this improvement attributable to sharpening of connectivity patterns irrespective of task condition (context-independent FC [ciFC]), task-evoked activation, and/or task context-dependent changes in FC (cdFC; Fig. 1a)? To explore this question, we modeled each of these signal components using psychophysiological interaction (PPI) analyses^22,23^, and tested their predictive utility using the connectome-based predictive modeling (CPM) framework^2,24,25^, with a focus on the most frequently studied terms: ciFC and activation. In a wide range of tasks in over 700 subjects, FC consistently predicted phenotype, while activation only predicted phenotype when the task was related to the predicted phenotype. Further, for each task, predictive FC and activation patterns were spatially distributed and relatively non-overlapping, with predictive ciFC concentrated in medial frontal and frontoparietal networks, and predictive cdFC concentrated in motor and visual networks; while all four networks have been previously implicated in predictive models of fluid intelligence^2,16^, this analysis framework is the first to permit insight into distinct, phenotyperelevant task effects on FC in these networks.

**Figure 1.**
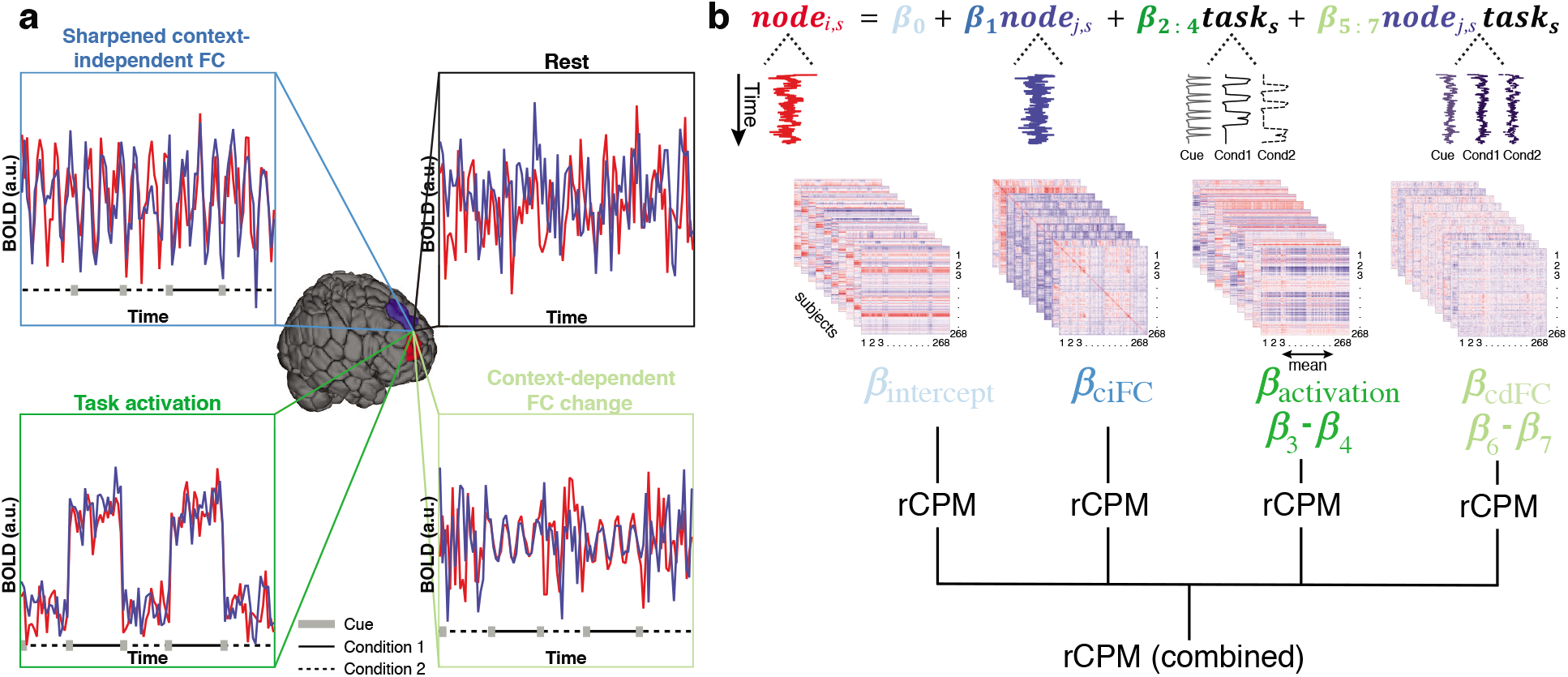
Pairing psychophysiological interaction (PPI) analysis with prediction permits characterization of how tasks change patterns of brain activity to reveal brain-phenotype relationships. (a) Relative to rest, in-scanner tasks may sharpen context-independent FC (“ciFC”), elicit baseline shifts in activity (“task activation”), and/or induce context-dependent changes in nodal synchrony (“cdFC”). (b) These components of each node’s time course (here, examples taken from nodes in the WM task; see Methods) can be modeled in an adapted PPI framework, yielding one node-by-node matrix of PPI betas for each term in each subject *s*. After calculating condition contrasts for the task activation and cdFC terms (indicated schematically by subtracted betas) and collapsing task activation via averaging into a single value per node per subject, these matrices (and vector, in the case of activation) were then submitted, individually and in combination, to the ridge CPM pipeline (rCPM)^25^ to yield predictions of fluid intelligence (gF). See Methods for details. Cond, condition; *i, j*, two nodes in the Shen parcellation^2,28^; *s*, subject.

Given this evidence that tasks are changing FC to improve prediction, we next sought to characterize these predictive changes. Motivated by the finding that tasks increase both the similarity of individuals’ FC and the identifiability of individuals on the basis of FC patterns^15^, we developed an extension of PPI and inter-subject FC^26^ analyses. Results demonstrate a nuanced relationship between FC predictive utility and inter-subject consistency: moderate, but not high, consistency aids prediction. Together, these findings suggest that tasks meaningfully change whole-brain functional organization, revealing brain-phenotype relationships by synchronizing individuals’ brains just enough to de-noise the BOLD signal, while also preserving—indeed, amplifying—meaningful individual differences in FC.

## Results

To explore why task FC-based models outperform resting-state FC-based models, we used data from the Human Connectome Project (HCP)^27^, S1200 release (*n* = 703; see Methods for inclusion criteria). Each subject completed seven in-scanner tasks, providing an internal validation of results’ generalizability; for each task run, fMRI data were parcellated into 268 nodes^2,28^ and a mean time course was calculated for each node. Each node’s time course was decomposed via multilinear regression, using a validated psychophysiological interaction (PPI) framework^22,23^, into terms that reflect its context-independent FC with the predictor node (ciFC), its context-dependent FC with the predictor node (cdFC), its task activation, and its overall degree of task-induced signal change (for further discussion of this term, see Supplementary Fig. 1 and Methods*: Investigating potential confounds)*. This regression was performed for every node pair for each task and subject, and linear contrasts were calculated for task activation and cdFC terms, yielding four PPI beta matrices per subject per task (Fig. 1b).

These matrices were then submitted, individually (“individual-term model”) and in combination (“combined model”), to a ridge regression-based version of the connectome-based predictive modeling pipeline (rCPM)^24,25^ to predict phenotype (here, gF) scores (Fig. 1b). In brief, this cross-validated machine learning approach selects features on the basis of their correlation with the predicted measure, regresses (with ridge regularization) phenotype scores on selected features’ values (here, PPI beta estimates), and uses resulting ridge regression coefficient estimates to construct a linear model relating brain data to phenotype measures. This model is then applied to the left-out fold, and the process is repeated iteratively until all folds have been used as the test group. Model performance was quantified as 1 – normalized mean squared error (Methods: *Cognitive prediction*) of each model (*q*^2^; higher values indicate better performance). This analysis was repeated 90 times with different assignments of participants to folds, and performance is presented for every iteration (Fig. 2a) and as the mean and standard deviation across iterations and tasks for each term (Fig. 2c).

**Figure 2.**
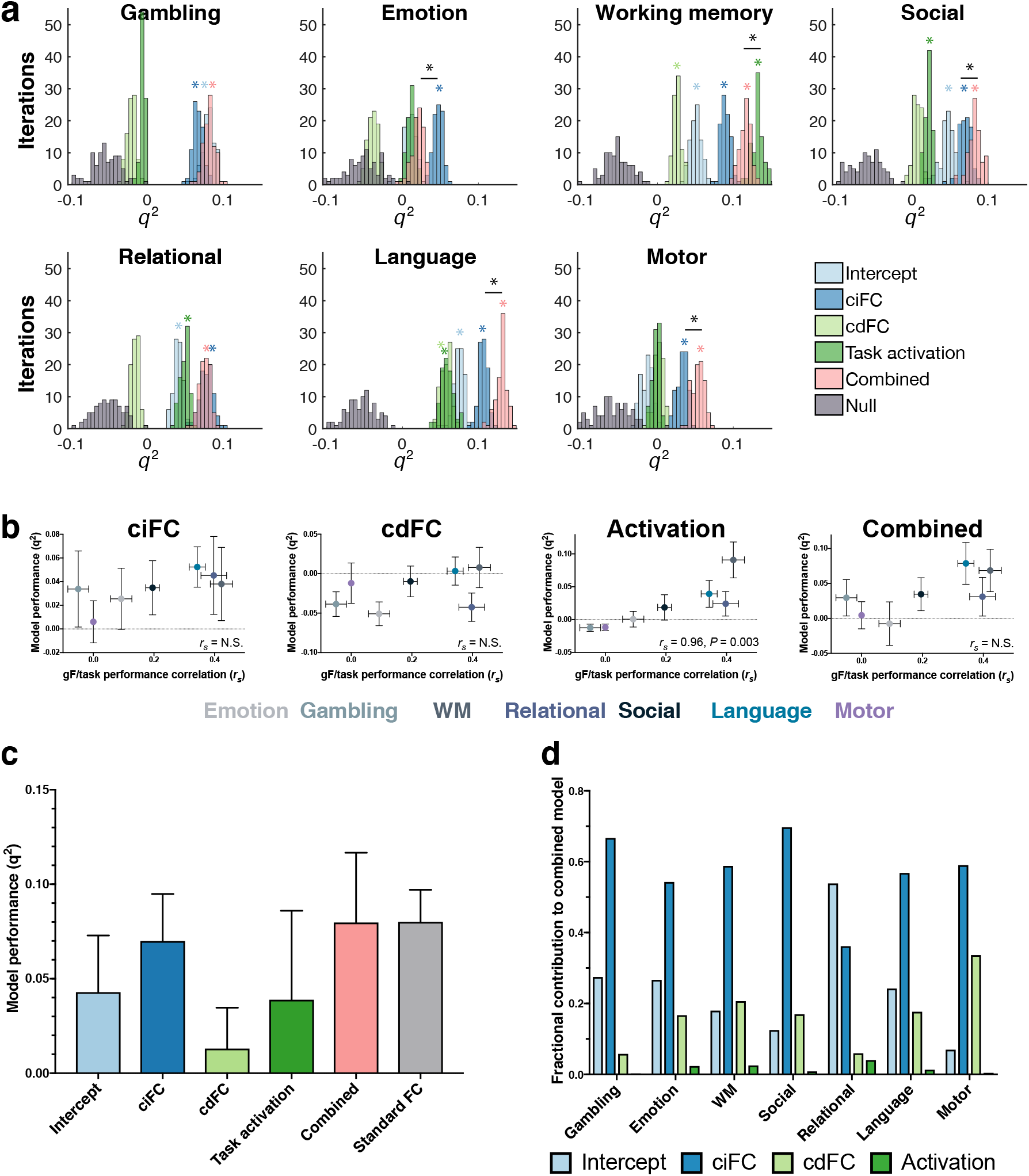
Context-independent FC, context-dependent FC, activation, and overall task effect contain complementary phenotypic information. (a) Histograms of model performance (quantified as *q^2^* [see Methods]) across 90 iterations of 10-fold rCPM with a *P* threshold of 0.1 for feature selection. Black asterisks, tasks for which the best-performing model significantly outperformed the second-best-performing model (\**P* < 0.001, corrected, via Wilcoxon signed rank test). Colored asterisks, terms that significantly predicted gF, via 90 iterations of nonparametric permutation tests. “Null”: prediction performance using permuted gF scores. (b) Mean task relatedness versus model performance for each model type. Error bars indicate standard deviation across 10 split-half analysis iterations (see Methods); *r_s_*, rank correlation between mean task relatedness and model performance across tasks. (c) Summary (mean and s.d., indicated by error bars) of model performance for each term, averaged across tasks. Models with negative mean *q^2^* were set to 0. “Standard FC”: prediction performance using FC calculated as Fisher-transformed Pearson correlations, without task modeling (see Methods). For a complete summary of all terms’ performance for each task separately, see Supplementary Fig. 3. (d) Percent contribution of each term to that task’s combined model (see Methods for derivation). WM, working memory.

Model performance was assessed for significance using non-parametric permutation tests. That is, the analysis was repeated 90 times with gF permuted across subjects each time; given the existence of many sibships in the dataset, allowed permutations respected family-related limits on exchangeability^29,30^. *P* values were calculated as the fraction of iterations on which the unpermuted gF-based models performed worse than or as well as the best-performing corresponding null model. For each task, the best-performing (i.e., highest mean *q*^2^) unpermuted gF-based model was also compared to the second best-performing model, and the combined model to the model based on “standard FC” matrices (i.e., FC calculated as Pearson correlation of node pairs’ time courses during a given task, without task modeling) via two-sided Wilcoxon signed-rank tests. All *P* values were corrected for multiple comparisons using the Bonferroni correction.

### Task-based FC predicts gF for all tasks, independent of activation

Consistent with previous work demonstrating that HCP task FC-based models successfully predict gF^16^, for each task, ciFC significantly predicted gF (all *P* < 0.01, corrected), regardless of whether or not task activation significantly predicted gF. In fact, ciFC yielded the most successful individual-term models across tasks (Fig. 2c). Only tasks that are most related to the predicted measure elicited predictive activation (Fig. 2b; task relatedness to gF [see Methods: *Cognitive prediction]* versus task activation-based prediction performance, *r*_s_ = 0.96, *P* < 0.05, corrected). No other model (ciFC, cdFC, or combined) demonstrated a significant relationship between task relatedness and prediction performance. Further, combining information from multiple signal components (combined model) further improved prediction performance (Fig. 2a, c): performance of the combined model was as good as (2 tasks; *P* > 0.05, corrected) or better than (3 tasks; *P* < 0.001, corrected) the best individual-term model in 5 out of 7 tasks, and better than models built from standard FC matrices in 2 tasks (*P* < 0.001, corrected). See Supplementary Table 1 for a comparison of these results with and without global signal regression.

Examining the relative contributions of each term to the combined models offers additional insight into term predictive utility (Fig. 2d; see Methods for derivation). These contributions often did not follow the performance of corresponding individual-term models for that task, highlighting the importance of information uniqueness. For example, cdFC alone did not significantly predict gF for the emotion, social, or motor tasks, but made substantial contributions to the combined models for each. This finding also held in reverse; that is, while model performance generally dropped more precipitously when high-contributing terms were dropped from them than when low-contributing terms were dropped, contribution did not entirely dictate performance (Supplementary Fig. 2). Notably, ciFC made the largest contribution to combined models for 6/7 tasks, while task activation made negligible contributions to combined models in any task (Fig. 2d).

Further, given that mean motion was significantly correlated with observed gF for 3/7 tasks (Supplementary Table 2) and with mean predicted gF for 9/35 tasks and terms (Supplementary Table 3), several additional analyses were performed to ensure that in-scanner motion did not confound results. First, rCPM was repeated using partial correlation with individuals’ mean motion per task—rather than simple correlation—for feature selection (i.e., selecting features that are correlated with gF after controlling for their correlation with motion). Second, even more conservatively, rCPM was repeated after regressing individuals’ mean motion for the given task from both gF and each edge (within the cross-validation loop); models were built from resulting residuals (Methods: *Investigating potential confounds*). Results are comparable, both in terms of mean model performance (*r*(original, partial correlation) = 0.99, *r*(original, residualized [10-fold]) = 0.99, all *P* < 0.001; Supplementary Table 4) and feature weights (Supplementary Table 5), suggesting that modeling results are not confounded by in-scanner motion. Frame-to-frame motion was also found to be uncorrelated with task timing (Supplementary Table 6), and repeating prediction analyses using only the half of the sample that moved the least (grand mean RMS relative motion < 0.072mm across tasks, *n*=351) also yielded comparable patterns of relative prediction performance (Supplementary Table 4).

### Model contributions of FC and activation terms are spatially distributed and distinct

We next sought to characterize the spatial distribution of features with high predictive utility (i.e., model contribution) for each term in each task’s combined model (see Methods for derivation). Because different insights can be gained from interpreting predictive contributions at the node, network, and edge levels^31^, we present results at all three levels of summarization. Except where otherwise noted, signed contributions were used to dissociate features that are positively and negatively related to gF. The seven tasks demonstrated substantial consistency in overall spatial patterns of predictive utility; for concision and clarity, the language task is used to exemplify these patterns, with corresponding results for all tasks displayed in Supplementary Fig. 4.

First, we explored the distribution of predictive features at the network level, grouping nodes into ten canonical networks defined in an independent sample^2,32^ (for details, see Methods and Supplementary Fig. 5). Results across tasks demonstrate the substantial and distributed contributions of ciFC (Fig. 3a, Supplementary Fig. 4a), cdFC (Fig. 3b, Supplementary Fig. 4c), and activation (Fig. 3f, Supplementary Fig. 4e) features to the combined models. Critically, predictive features are relatively non-overlapping for ciFC and cdFC (paired, two-sided Wilcoxon signed rank test of ciFC versus cdFC concatenated, vectorized, network matrices: *P* < 0.001; mean rank correlation between vectorized, absolute model contributions for ciFC versus cdFC across all tasks: overall 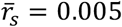, union of predictive ciFC and cdFC features only 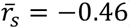). In particular, visual inspection of the difference between ciFC and cdFC model contributions— at both the network and node levels—reveals that, across tasks, predictive ciFC edges tend to be concentrated in medial frontal and frontoparietal networks, while predictive cdFC edges tend to be concentrated in motor and visual networks (average difference, Fig. 3c; difference for each task, Supplementary Fig. 4f).

**Figure 3.**
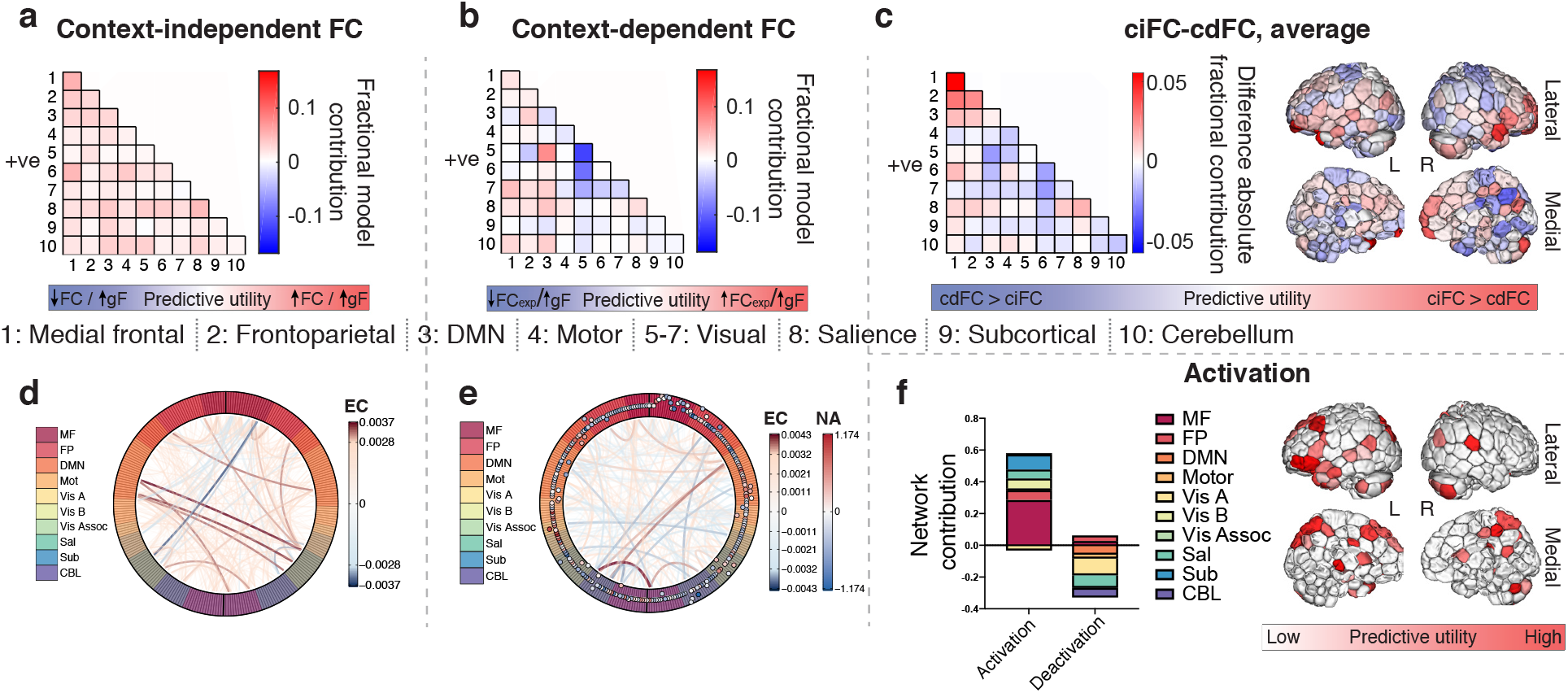
Context-independent FC, context-dependent FC, and activation predictive features are distributed and distinct. (a) Visualization of predictive ciFC features by network for the language task. Red = positive mean ridge coefficients, blue = negative mean ridge coefficients, shade = relative model contribution. In this and all subsequent figures: “+ve” indicates that results reflect only contributions of edges with mean positive ciFC. 1-10 = network assignment. (b) Visualization of predictive cdFC features by network for the language task. FC_exp_, FC during the experimental condition (i.e., condition of interest). Red = positive mean ridge coefficients, blue = negative mean ridge coefficients, shade = relative model contribution. 1-10 = network assignment. (c) Visualization of the differences in localization of predictive ciFC and cdFC. Left: difference between absolute contribution-based, network-level ciFC model contributions and absolute contribution-based, network-level cdFC model contributions, averaged across tasks. Right: difference between absolute node-level contributions (degree) of ciFC and cdFC (Methods). For both visualizations, blue indicates networks and regions with cdFC > ciFC predictive utility; red, ciFC > cdFC. (d) Visualization of individual predictive ciFC features for the language task, with each consistently selected edge represented as a line; line color and thickness scale with predictive model contribution. In this and all subsequent figures, MF = medial frontal, FP = frontoparietal, DMN = default mode network, Mot = motor, Vis A = visual A, Vis B = visual B, Vis Assoc = visual association, Sal = salience, Sub = subcortical, CBL = cerebellum. EC, edge contribution. (e) Visualization of individual predictive cdFC (lines) and activation (outer track circles) features for the language task; line color and thickness scale with cdFC feature predictive utility, and circles represent the corresponding nodes, with their color indicating mean activation (red = positive, blue = negative) and distance from the x axis indicating their model contribution. EC, edge contribution; NA, node activation. (f) Visualization of predictive activation features at the node and network levels for the language task. Left: fractional mean network contributions for predictive nodes with mean positive PPI activation betas (“activation”) and for nodes with mean negative PPI activation betas (“deactivation”). Right: absolute, fractional model contribution of nodes’ activation. Comparable results for all tasks are displayed in Supplementary Fig. 4.

It is important to note that the predictive utility and distribution of useful context-dependent features will be affected by the condition contrast that is applied, and these results demonstrate how findings can be interpreted with some cognitive specificity, given this choice. For example, while motor task ciFC and cdFC demonstrate similar overall patterns of predictive utility as other tasks (i.e., relative overrepresentation of edges in medial frontal and frontoparietal networks for ciFC, and in visual networks for cdFC), the motor task contrast—all motions (tongue, hands, and feet) versus fixation—reveals consistently negative cdFC ridge coefficients. That is, across the brain, the lower the FC during these body motions, the higher an individual’s gF (Supplementary Fig. 4c). This pattern is in stark contrast to other tasks’ patterns of cdFC predictive contributions, a finding with potentially interesting cognitive implications (see Discussion).

Before calculating the predictive contributions of network pairs (or networks, in the case of activation), ciFC and cdFC features were divided into edges with mean positive and mean negative ciFC (and activation features into nodes with mean positive and negative activation; see Methods: *Evaluating and visualizing contributions to a predictive model)*. That is, we sought to approximately separate edges that have, at baseline, positive FC (mean positive ciFC) from those that have, at baseline, negative FC (mean negative ciFC). We found that ciFC and cdFC of negative edges made essentially no contributions to combined models for any of the tasks (Supplementary Fig. 6a-b); we note, however, that there were substantially fewer negative than positive edges (overall and selected), likely contributing to this result. For these reasons, main results reflect only positive edges (Fig. 3a-c, Supplementary Fig. 4a, c, f). Network-level contributions of predictive cdFC patterns were further divided by mean cdFC (i.e., edges that strengthen during the experimental condition [positive mean cdFC] and edges that weaken during the experimental condition [negative mean cdFC]); results are visualized in Supplementary Fig. 7. Conversely, both activated and deactivated nodes were consistently included in predictive models; given this, task activation network-level predictive utility is presented for both activated and deactivated nodes (Fig. 3f; Supplementary Fig. 4e).

Visualizing predictive contributions at the network level, while useful, offers a necessarily coarse representation of where in the brain highly predictive features are located. To visualize the predictive utility of individual features, we present circle plots, with each network a different color on the outer track, and each line a ciFC (Fig. 3d, Supplementary Fig. 4b) or cdFC (Fig. 3e, Supplementary Fig. 4d) edge. This visualization again highlights that model contributions are distributed and relatively non-overlapping across terms. Specifically, it reveals that highly predictive cdFC features are not incident to the most predictive or most activated nodes (Fig. 3e, Supplementary Fig. 4d): mean correlation of cdFC absolute node degree (Methods) with absolute predictive utility of node activation: 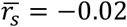; mean correlation of cdFC absolute node degree with absolute task effect size: 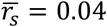. Similarly, the absolute contribution of a node’s activation is only weakly related to its absolute task effect size: 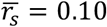. In sum, these results demonstrate that predictive utility—of ciFC, cdFC, or even task activation, itself—is not simply driven by task activation, and that predictive features from each of these terms are spatially distributed and relatively non-overlapping.

### Context-independent FC is more consistent across individuals than context-dependent FC or activation

To explore the consistency of task effects on FC and activation across individuals, and the potential relationship between this consistency and predictive utility, we performed an inter-subject PPI analysis (Fig. 6; Methods: *Inter-subject psychophysiological interaction analysis*) on the five tasks with consistent task timing across individuals (emotion, gambling, social, relational, and WM).

This analysis revealed substantial consistency in activity patterns across individuals, highlighting two, task-general trends. First, during a task, positive edges’ ciFC and nodes’ time courses become more similar across individuals (relative to intrinsic components of the BOLD signal, e.g., at rest, which should be uncorrelated across subjects^26^) in almost all networks for all tasks (Fig. 4a). On the other hand, context-dependent effects vary more by task and network, as would be expected given the varying designs and demands of the tasks (e.g., some network pairs’ edges were consistently stronger during the experimental condition relative to the control condition [red], while others were consistently weaker during the experimental condition relative to the control condition [blue]; Fig. 4b). Second, ciFC inter-subject consistency was overall greater than cdFC or task activation inter-subject consistency (Fig. 4c; median ciFC consistency: mean across tasks = 0.396, range = 0.27-0.63; median cdFC consistency: mean across tasks = −0.01, range = −0.04-0.01; median activation consistency: mean across tasks = −0.004, range = −0.03-0.01), suggesting that moment-to-moment fluctuations are more similar across individuals than are block-level changes in FC and activation.

**Figure 4.**
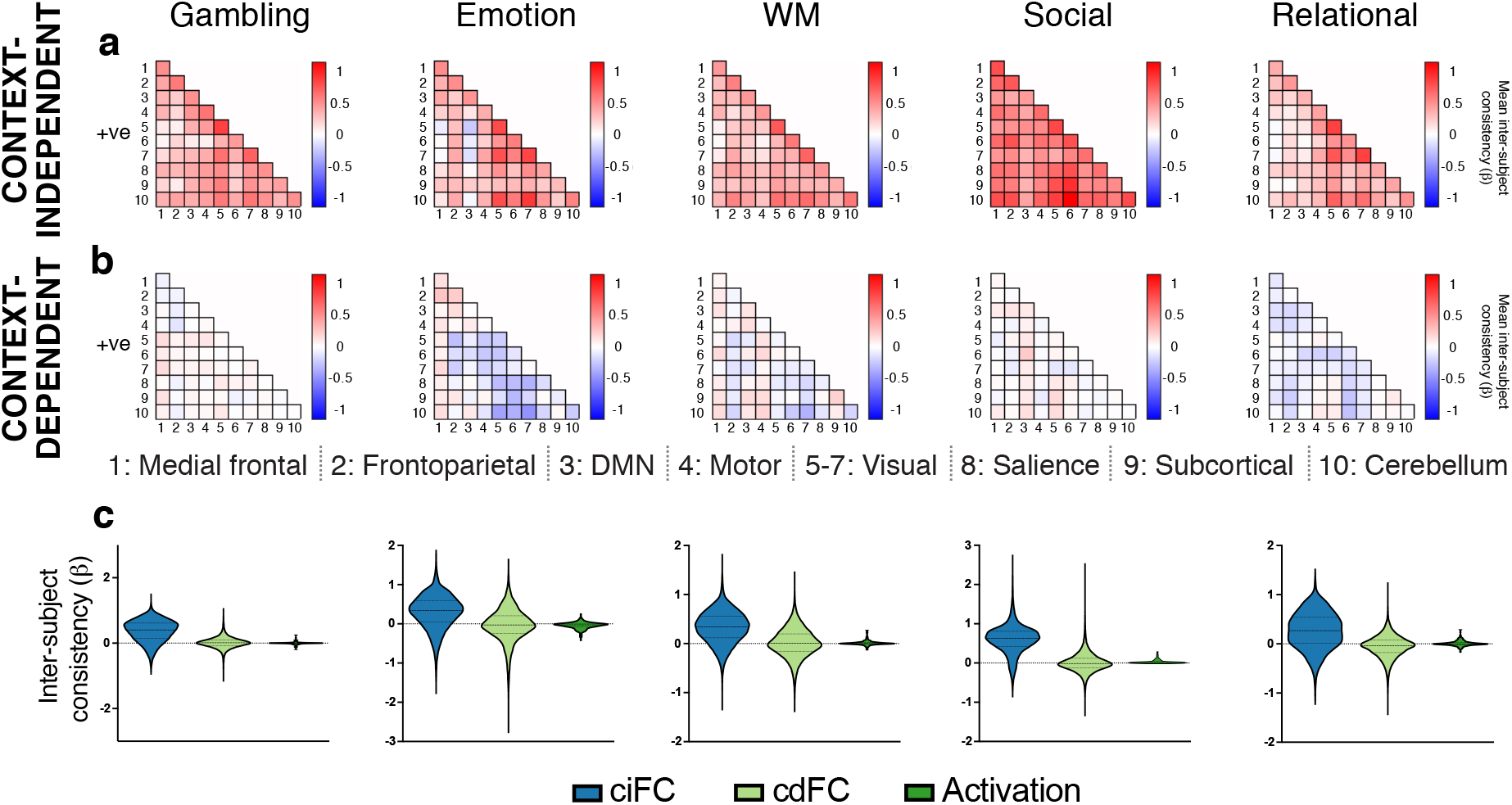
Inter-subject PPI analysis reveals consistent task-induced changes in context-independent and context-dependent activity across individuals. (a,b) Network-level visualization of the substantial inter-subject consistency for both context-independent and context-dependent signals. 1-10 = network assignment. (c) Violin plots (dashed line, median; dotted line, quartiles) of inter-subject consistency for all unique features (i.e., inter-subject PPI betas) reveal that inter-subject consistency of moment-to-moment fluctuations (ciFC) is greater than inter-subject consistency of block-level activation (“Activation”) or FC (cdFC) changes.

As was the case for the prediction analyses, effects were almost entirely limited to edges with positive mean ciFC (i.e., positive mean PPI betas for the ciFC term, as were used previously); inter-subject consistency of mean negative edges’ ciFC are presented in Supplementary Fig. 6c, and of mean negative edges’ cdFC in Supplementary Fig. 6d.

### Tasks increase predictive utility by increasing consistency—to a point

Given the substantial inter-subject consistency in ciFC, we next sought to explore whether there is any relationship between this consistency and predictive utility. To do so, we first evaluated the inter-subject consistency of reliably predictive and non-predictive features (Fig. 5a). In all five tasks, predictive edges were significantly more consistent across individuals than non-predictive edges (all *P* < 0.001, corrected, via two-sided Wilcoxon rank sum test; *g* = 0.11-0.64, mean = 0.34, via Hedges’ g), suggesting that edges that are modulated by the task are more useful for prediction.

**Figure 5.**
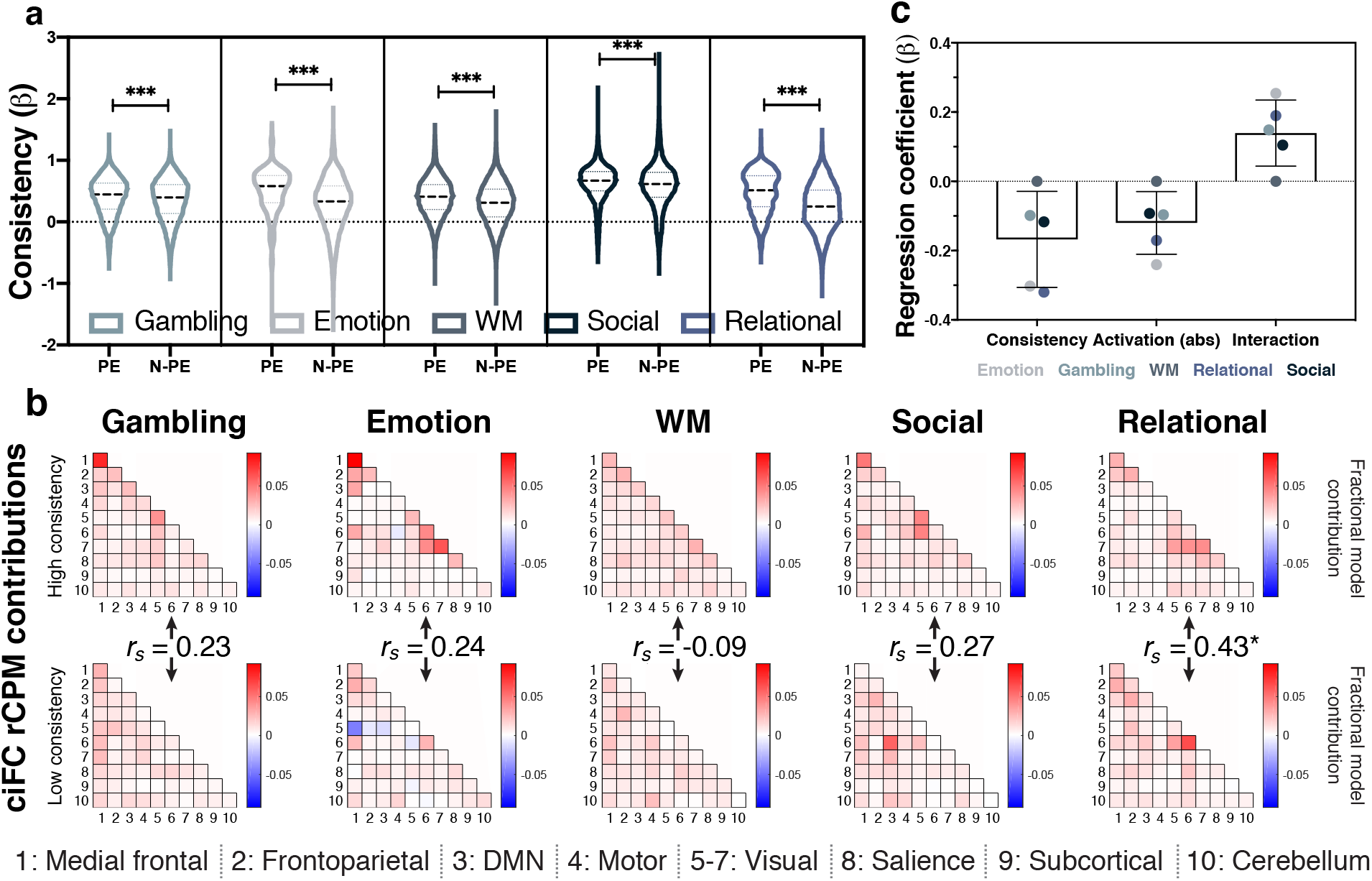
Predictive edges tend to be task modulated, with this modulation revealing signal components that meaningfully vary across subjects. (a) Violin plots (dashed line, median; dotted line, quartiles) of ciFC inter-subject consistency for reliably predictive edges (PE) and non-predictive edges (N-PE). \*\*\**P* < 0.001, corrected, Wilcoxon rank sum test. (b) Visualization of predictive ciFC features at the network level, with features divided by median, absolute inter-subject consistency. *r_s_*, rank correlation between high- and low-consistency network matrices for the given task and term, \**P* < 0.05, corrected. 1-10 = network assignment. (c) Regression analyses relating the predictive utility of a feature’s ciFC to its inter-subject consistency, task activation, and their interaction. Results presented as regression coefficient for the given predictor in each of the five modeled tasks; bar height reflects mean coefficient across tasks; error bars indicate s.d. of coefficients.

Next, in two, related analyses, we asked whether, within this set of predictive edges, predictive utility is related to the magnitude of inter-subject consistency. First, to identify any differences in the spatial localization of high- and low-consistency predictive edges, we divided reliably selected ciFC predictive features into high- and low-absolute inter-subject consistency groups. CiFC contributions were found to be spatially distinct for high- and low-consistency edges; this visual trend was confirmed by the low correlation between vectorized high- and low-consistency predictive utility network matrices: *r_s_* = −0.09 – 0.43, mean *r_s_* = 0.22, 4/5 tasks: *P* > 0.05, relational task: *P* < 0.05, all Bonferroni corrected (Fig. 5b).

Second, we explicitly modeled the relationship between absolute inter-subject consistency and absolute predictive utility using a multilinear regression: predictive utility of reliably predictive edges was regressed on inter-subject consistency, one additional anatomical or functional term and the interaction between this term and inter-subject consistency (Methods: *Modeling the relationship between inter-subject consistency and predictive utility*). To capture key anatomical and functional features for each edge that might moderate the relationship between consistency and predictive utility, we used the following terms as predictors of predictive utility: mean absolute task effect size for the two nodes incident to the given edge (calculated as in ^33^), edge membership between or within canonical networks, resting-state edge test-retest reliability (calculated as in ^32^), edge length (i.e., Euclidean distance between incident nodes), and edge location within or between hemispheres.

Overall, inter-subject consistency was negatively related to predictive utility, such that more consistent edges were less predictive. Interestingly, activation was also negatively related to predictive utility, suggesting that activated regions are connected by less predictive edges than non-activated regions. The interaction between inter-subject consistency and activation, however, was positive (Fig. 5c), suggesting that edges affected by task activity are more predictive when that effect is consistent across individuals. Hemisphere, edge length, network, and reliability were not clearly related to predictive utility; results for these analyses are displayed in Supplementary Fig. 8.

## Discussion

In-scanner tasks have been found to amplify individual differences in patterns of FC and correspondingly improve FC-based prediction of phenotype^16^, but the nature of this improvement—whether (and how) it is due to changes in ciFC, cdFC, and/or activation patterns—remains unexplored. In this work, we leverage intra- and inter-subject PPI and predictive modeling analyses to ask this question. Despite substantial differences in the nature and design of the analyzed tasks, we found a striking consensus: distributed, task-induced changes in FC predict phenotype independent of task activation. That ciFC successfully predicts phenotype for all tasks and is, across tasks, the best-performing individual-term model suggests that FC, not activation, drives improved prediction performance of FC-based models that utilize task data. Further, these task-induced changes in FC best reveal brain-phenotype relationships when they are moderately consistent—but not too consistent—across individuals.

Together, these results demonstrate that tasks have widespread effects on patterns of brain activity, of which focal task activations are only a small part, and that brain-phenotype relationships are best revealed by measures, such as FC, that capture these distributed effects. These findings are in line with the growing consensus that complex cognitive processes and constructs, such as fluid intelligence, are supported by distributed neural circuitry, rather than by circumscribed regions of interest^34^, and demonstrate that rest-to-task FC changes, while perhaps small in magnitude^9^, contain important information about phenotype independent of task-evoked activation. We of course do not suggest that co-activation cannot drive changes in FC (see, for example, ^17^), but rather that FC changes relevant to predictive models are not driven by co-activation.

### Anatomy of a successful predictive model

It is noteworthy that the components of a given node’s signal (both FC and activation) contain complementary (i.e., unique), phenotype-relevant information, such that their combination in many cases further improves predictive model performance for a given task; in fact, in many cases, individual terms contribute more to the combined model than would be expected given the performance of corresponding individual-term models. These findings are consistent with reports of task activation and FC reflecting unique information about in-scanner task performance^19^, and of improved ridge regression-based CPM performance with the inclusion of more relevant features in the model (up to a *P* value of 0.5)^25^. That is, in evaluating and weighting each component of the signal separately, the combined model is able to capture information that is not accessible from its component parts. Efforts to reveal brain-phenotype relationships using regularized regression-based models may therefore benefit from the inclusion of more features, even if their relationship to the phenotype of interest is relatively weak^25^, although it is always wise to exclude uninformative features to avoid overfitting^35^.

Interrogation of these combined models revealed that they are broadly distributed and relatively non-overlapping across terms (i.e., ciFC, cdFC, and activation); that is, predictive ciFC and cdFC edges were distinct, and predictive edges were not incident to nodes with the most predictive activation. This further highlights the distinct, phenotype-relevant ways in which tasks alter each signal component and replicates the finding that task-related activation and context-dependent FC are spatially distinct^18^. Further, across all tasks, predictive FC patterns tend to be concentrated in medial frontal, frontoparietal, visual, and motor networks. These networks have all been previously implicated in successful FC-based predictive models of gF^2,16^, but PPI-based prediction permits a more fine-grained evaluation of their involvement, revealing that medial frontal and frontoparietal networks are overrepresented in predictive *context-independent* FC edges, while visual and motor networks are overrepresented in predictive *context-dependent* FC edges (Fig. 3c, Supplementary Fig. 4f). That is, while medial frontal and frontoparietal network FC is relevant to gF regardless of when in the task you look, block-level changes in visual and motor network FC predict gF. This is consistent with evidence that frontal and frontoparietal networks comprise domain-general, core components of task-set representations^23,36^, while visual and motor networks comprise domain-specific “data processing” systems^37^, which would be expected to adapt their operations to the nature and demands of the task at hand.

This work is, to our knowledge, the first to parse and localize the differential predictive utility of context-independent FC and context-dependent FC. Results demonstrate the exciting potential of this analysis framework to understand the neural bases of successful predictive models, offering more nuanced insights into the neural representation of the predicted measure than would be accessible with standard FC-based models.

### A framework to explore task-specific effects on functional organization

In addition to task-general predictive changes in FC, spatial localization of predictive features demonstrates the utility of this approach for drilling down into task-specific FC changes and their relationships to phenotypic measures. For example, the motor task is the only task for which cdFC model contributions at the network level are consistently negative, indicating that the weaker an individual’s predictive cdFC edges during motion, the higher that individual’s gF. Further, the finding that both edges that strengthen (i.e., mean positive cdFC) and those that weaken (mean negative cdFC) during motion relative to fixation have negative context-dependent FC ridge coefficients (Supplementary Fig. 7) suggests that the relationship between cdFC and gF depends more on total FC strength (i.e., globally weaker FC during motion predicts higher gF) than on the nature of task-induced change in FC (i.e., increased or decreased edge strength). It is possible that this task, given its low cognitive demands, does not require the widespread neural interactions that support higher-demand tasks^38^, permitting mind wandering and decreased neural integration, which is energetically costly^18,39^.

While these results would be expected to depend to some extent on task design and modeling choices^40^, they suggest a potentially fruitful direction for future investigation into the nature and cognitive implications of task-specific changes in functional brain organization. The utility of such task-specific changes to predict performance on that task has been demonstrated for an inhibitory control task^19^; here, we broaden this idea to suggest that task-specific changes in brain activity may improve prediction of less directly related phenotypes. It is likely that such task-specific changes, by offering complementary insights into brain-phenotype relationships, explain the finding that combining FC data across task conditions often outperforms prediction using FC data from a single condition^25,41^. Better understanding these changes will enable more selective inclusion of data by condition, particularly when using data that include potentially less informative or noisy conditions (e.g., rest^41^).

### Inter-subject PPI reveals how tasks increase FC predictive utility

The inter-subject PPI analyses are a novel extension of prior work on inter-subject correlation^42^ and FC^26^ that provide a complementary approach to study how tasks change patterns of FC to better reveal meaningful individual differences in them. Given the findings that tasks both increase inter-subject FC similarity and improve individual identifiability on the basis of FC patterns^15^, and that moment-to-moment “events” in the BOLD signal explain substantial FC variance, are highly synchronized across individuals during a task, and are time points at which individual identifiability is maximized^43^, we sought to explicitly explore the relationship between inter-subject time course synchrony (i.e., consistency) and predictive utility: do tasks constrain the state space^44–46^, simultaneously making a relevant network more similar across individuals and amplifying signal components within it that vary across individuals, and/or are increased inter-subject consistency and increased predictive utility spatially separable? In support of the former, predictive edges have higher inter-subject consistency than non-predictive edges. Among these predictive edges, however, predictive utility is highest in edges that are least similar across individuals. Relatedly, among predictive edges, activation of incident nodes decreases an edge’s predictive utility overall, but activated edges are most useful when activation patterns are consistent across individuals. In both cases, some consistent task modulation reveals FC patterns that set individuals apart. In line with these findings, high-consistency predictive edges were more concentrated in frontal and visual networks, which are also the networks most commonly activated by these tasks.

Taken together, these results suggest that tasks reveal FC-phenotype relationships by making edges moderately consistent—but not too consistent—across individuals. This finding may be explained by two, related task effects: (1) Tasks synchronize individuals’ brain activity by time locking it to the task^42^ and thus making it more constrained^44^. This may decrease edges’ “noisy” sources of inter-subject variability that dominate the resting state^15^, which would increase their inter-subject similarity *and* their predictive utility. Some between-subject variance, however, reflects meaningful (i.e., phenotypically relevant) inter-individual differences in functional brain organization^2,32^; eliminating this variance would further increase edges’ inter-subject consistency but *decrease* their predictive utility. Tasks may therefore improve FC-based prediction performance by de-noising the BOLD signal, revealing, through this moderate synchrony, meaningful individual differences in FC.

### Additional considerations and future directions

A deeper investigation of task-specific activation patterns may reveal that task-evoked activation here demonstrates little predictive utility (c.f., ^47^) because of the relatively large size of each node in our selected atlas^2,28^, which may blur informative, fine-scale patterns of activity and/or “wash out” activated voxels by grouping them with less activated voxels^34,48,49^. Crucially, we do not seek to make claims about the predictive utility of task activation, but rather to demonstrate that task-induced changes in standard FC that improve phenotype prediction are not driven by task activation. Given this, along with previous demonstrations that number (and thus size) of nodes does not affect prediction performance^16^, we chose to use a conventional parcellation to calculate FC^2,28^. However, future work may seek to compare the predictive utility of FC and task activation at a finer spatial scale or using task-specific parcellations, given the recent demonstration that node boundaries meaningfully reconfigure with task^33^. Our finding that activation is predictive only for tasks that are relevant to the predicted phenotype, consistent with the recent demonstration that activation evoked by more cognitively demanding tasks best predicts general cognitive ability^47^, suggests that such work may be particularly fruitful in the case of task-phenotype correspondence.

Similarly, our characterization of task effects is intentionally limited. Given that blocked task designs are common and better powered to reveal effects of interest than event-related or mixed designs^50,51^, we chose to use simple, well-studied task condition contrasts^52^ to reveal fundamental, generalizable effects of task execution on FC and activation predictive utility. Finally, regression analyses are here limited by the relatively small number of measurements (i.e., coefficient estimates for the five tasks that were used for both inter- and intra-subject analyses). Future application of the analysis framework presented here to independent datasets will provide opportunities to replicate and broaden presented results.

In particular, as the human neuroimaging community increasingly uses task FC, rather than resting-state FC, for phenotypic prediction, our understanding of how best to do so will be furthered by asking “which tasks for what phenotype?”. Our results suggest that the optimal task for prediction is one that synchronizes individuals’ brain activity just enough to de-noise the BOLD signal, but not so much that meaningful individual differences in brain functional organization are obscured. Efforts to optimize these criteria, perhaps using a continuous or naturalistic paradigm such as movie viewing^53^, and to expand them by identifying relevant, task-specific effects present exciting opportunities for future work.

### Conclusion

As task-based FC proves its utility to reveal brain-phenotype relationships, a better understanding of how tasks change patterns of FC is critical. By demonstrating that the success of task FC-based predictive models is attributable to task-induced changes in FC—regardless of task activation—and by characterizing how tasks change patterns of context-independent FC to improve prediction, these findings demonstrate that reconfiguration of the functional connectome during in-scanner tasks is real, meaningful, and useful. This lays the foundation for intentional, precise use of in-scanner tasks to amplify individual differences in functional brain organization in meaningful ways and more effectively map the neural representations of behaviors, traits, and clinical symptoms.

## Materials and methods

### Dataset

Data used in this work were released as part of the Human Connectome Project (HCP) S1200 release, described below.

#### HCP participants

We restricted our analyses to those subjects who completed all seven fMRI tasks (WM, gambling, language, social, relational, motor, and emotion), whose grand mean root mean square (RMS) relative motion across all task runs was less than 0.1 mm and whose maximum mean RMS relative motion was less than 0.16 mm, and for whom gF measures were available. One subject was found to be missing data due to a download failure, and was excluded from all analyses to ensure consistency with previous results. A similarly conservative threshold for motion-based exclusion has been previously demonstrated to mitigate the relationship between FC and gF measures^16^. In total, data from 703 subjects were used (342 males, ages 22-37 years [mean = 28.5, s.d. = 3.8, median = 29]).

#### HCP imaging parameters and preprocessing

Details of imaging parameters^27,54,55^ and preprocessing^55,56^ have been published elsewhere. In brief, all fMRI data were acquired on a 3T Siemens Skyra using a slice-accelerated, multiband, gradient-echo, echo planar imaging (EPI) sequence (TR = 720 ms, TE = 33.1 ms, flip angle = 52 degrees, resolution = 2.0 mm^3^, multiband factor = 8). Images acquired for each subject include a structural scan and eighteen fMRI scans (WM task, incentive processing [gambling] task, motor task, language processing task, social cognition task, relational processing task, emotion processing task, and two resting-state scans; two runs per condition [one left/right (LR) phase encoding run and one right/left (RL) phase encoding run]), split between two sessions^52,55^. Data from the seven HCP tasks were used for this work, and each task was a different length (WM, 5:01; gambling, 3:12; language, 3:57; social, 3:27; relational, 2:56; motor, 3:34; emotion, 2:16). The scanning protocol (as well as procedures for obtaining informed consent from all participants) was approved by the Institutional Review Board at Washington University in St. Louis. Use of HCP data for these analyses was deemed exempt from IRB review by the Yale Human Investigation Committee. The HCP minimal preprocessing pipeline was used on these data^56^, which includes artifact removal, motion correction, and registration to standard space. All subsequent preprocessing was performed in BioImage Suite^57^ and included standard preprocessing procedures^2^, including removal of motion-related components of the signal; regression of mean time courses in white matter and cerebrospinal fluid; removal of the linear trend; and temporal smoothing with a Gaussian filter, σ = 0.18 (a relatively high low-pass filter designed to preserve potential high-frequency, task-related signal components). Mean RMS relative motion was averaged for the LR and RL runs, yielding seven motion values per subject; these were used for subject exclusion and motion analyses (e.g., partial correlation-based feature selection). All subsequent analyses and visualizations were performed in BioImage Suite^57^, Matlab (Mathworks), R version 3.6.0 for macOS (packages: RColorBrewer^58^, ComplexHeatmap^59^, and circlize^60^), and GraphPad Prism version 8.0 for macOS.

### Functional parcellation and network definition

The Shen 268-node atlas^2,28^ was applied to the HCP data, as described previously^16^. This parcellation is derived from the application of a group-wise spectral clustering algorithm to an independent data set^28^. Time courses of voxels within each node were averaged. Subjects without whole-brain coverage (i.e., with missing nodes) were excluded from all further analyses.

The same spectral clustering algorithm was used to assign these 268 nodes to eight networks^2,28^, and the subcortical-cerebellar network was split into networks 8-10 (Supplementary Fig. 5)^32^. These networks are named based on their approximate correspondence to previously defined resting-state networks, and are numbered as follows: 1. Medial frontal, 2. Frontoparietal, 3. Default mode, 4. Motor, 5. Visual A, 6. Visual B, 7. Visual association, 8. Salience, 9. Subcortical, 10. Cerebellum.

### Psychophysiological interaction (PPI) analysis

After parcellation, node time courses were submitted to an adaptation of the PPI pipeline developed and described by Cole and colleagues^23^ and modeled after the generalized PPI framework^22^. In brief, the mean time course for each node was decomposed via multilinear regression with three regressor types: the mean time course of a predictor node (yielding a beta weight that reflects context-independent FC [ciFC] between the predictor and target nodes); zero-centered, block-level task boxcar regressors convolved with the canonical HRF (yielding a beta weight that reflects the influence of task activation on the target node time course), and the interactions of these terms (yielding a beta weight that reflects context-dependent FC [cdFC] between the predictor and target nodes). The canonical HRF (generated using SPM8) was used given its demonstrated efficacy in identifying patterns of task activation in these data^52^. Task conditions and cues were modeled separately, as relevant (WM: 0-back task, 2-back task, and cue regressors; language: story task and cue regressors; emotion: face block and cue regressors; gambling: reward block and loss block regressors; relational: relational block, matching block, and cue regressors; social: mental video block and random video block regressors; motor: left hand, right hand, left foot, right foot, tongue, and cue regressors). Regressors were calculated separately for each subject using HCP E-Prime and EV .txt files and downsampled given the HCP sampling rate (i.e., TR = 720 msec). Thus, for a given task with *k* conditions (including fixation and cue, if present), a given node’s time course can be described by the following equation:

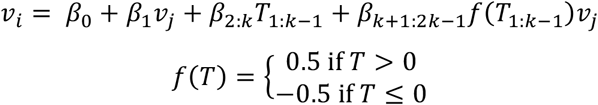

where *v_i,j_* are the *z*-scored time courses of target node *i* and predictor node *j*, and *T* is the relevant, HRF-convolved, zero-centered task timing regressor(s). As in previously published work^23^, the task regressors used in the interaction term were binarized. Here, after convolution with the HRF, values greater than zero were set to 0.5, and values less than or equal to zero were set to −0.5 (to zero-center the binarized task regressors, as for the non-binarized task regressors) prior to their multiplication by the predictor node time course to yield the interaction regressor. (For a discussion of the choice to zero-center task regressors, see Methods*: Investigating potential confounds*). These steps are each depicted schematically in Supplementary Fig. 9. For all tasks except the emotion and language tasks (for which there were no fixation blocks, rendering the contrast implicit by modeling only one of the two task conditions) and the motor task (for which all motion conditions were summed to yield a motion versus fixation contrast), interaction and task activation beta weights were each combined via subtraction to yield one activation contrast beta weight and one interaction contrast beta weight per feature in each subject^22^ (contrasts: *n*-back: 2-back – 0-back; gambling: reward – punish; emotion: fear [faces] versus neutral [shapes]; language: story versus math; relational: relation – match; social: TOM – random), and beta weights were calculated separately for each task run (i.e., LR and RL phase encoding runs) and then averaged. This process was repeated for every node pair (i.e., for a given target node, all other nodes were used as predictor nodes) and every subject, yielding, for each subject, four asymmetric, node-by-node matrices of beta weights (one each for the intercept, ciFC, task activation, and cdFC terms). Each task activation matrix was collapsed via averaging into a 268-element vector. All other matrices were symmetrized by averaging them with their transpose. All matrices (and vectors, in the case of activation) of a given type were then submitted—alone and in combination—to the predictive modeling pipeline described below.

### Cognitive prediction

Fluid intelligence was quantified using a 24-item version of the Penn Progressive Matrices test; this test is an abbreviated form of Raven’s standard progressive matrices^61^. Integer scores indicate number of correct responses (PMAT24_A_CR, range = 5-24, mean = 17.70, s.d. = 4.43, median = 19).

A modified version of connectome-based predictive modeling (CPM)^2,24^ was used to predict gF from brain measures (i.e., beta matrices [see *Psychophysiological interaction analysis])* using ridge regression^25^. This pipeline predicts gF in novel subjects, validating the model through iterative, *k*-fold cross-validation; in this work, *k* = 10 to balance model bias and variance given the large sample size^35^. Consistent with this motivation, split-half (i.e., *k* = 2) analyses yielded comparable patterns of results, but overall lower prediction performance (Supplementary Table 4). First, the sample was divided into ten groups, respecting family structure such that family members were always assigned to the same group. Nine of these groups were used as training data; in this training set, features (edges and/or nodes) were selected on the basis of their Pearson correlation with gF scores. A correlation *P* value of 0.1 was selected as the edge selection threshold, given evidence to suggest that more permissive feature selection yields improved regularized regression-based prediction results, and that *P* = 0.1 offers an acceptable compromise between model performance and computational demands^25^. These edges were then submitted as predictors (with gF score as response) to an L2-constrained linear least squares regression (elastic net mixing parameter = 1e-9). The regularization strength (λ) for this regression was optimized using another, inner 5-fold cross-validation, with λ set to the largest value that yields a MSE within one standard error of the minimum MSE. λ_max_ was computed using elastic net mixing parameter = 0.01 to find λ_max_ that shrinks all coefficients to 0 (which is impossible in the pure ridge case). This inner-loop hyperparameter selection implemented a separate feature selection on all available edges (i.e., not those remaining after the outer-loop feature selection) to avoid circularity. These fitted coefficients were then applied to the corresponding edges in the left-out test subjects to predict their phenotype scores, and these steps were performed iteratively with each group left out once.

Model performance was quantified as cross-validated 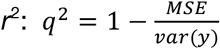, where *y* = observed gF scores (results were comparable using the correlation between observed and predicted gF as a performance metric; Supplementary Table 7). This whole pipeline was repeated 90 times with different group partitions, and model performance is reported as the mean across these 90 iterations (similarly, in graphical representations of results, bar heights represent mean performance, and error bars represent performance standard deviation). Significance of model performance was assessed via 90 iterations of nonparametric permutation testing^2^, accounting for limits on exchangeability due to family structure^29,30^, and *P* values were calculated as the fraction of non-permuted iterations on which prediction accuracy was less than or equal to the accuracy of the best-performing null model for the given task and term. Two related subjects in our sample were missing family structure information; these subjects were excluded from permutation tests (*n* = 701). Resulting *P* values were Bonferroni corrected for multiple comparisons. Last, for each task, paired Wilcoxon signed rank tests were used to compare performance across all 90 iterations of the two models with the highest and second-highest mean performance, as well as between the combined and standard FC-based models.

To compare the relatedness of the in-scanner task to gF to the utility of brain measures acquired during that task to prediction of gF, we performed an additional analysis. To avoid circularity, in one half of the sample, we performed 10-fold, cross-validated gF prediction, as before. In the second half, we rank correlated task performance (emotion: overall accuracy [variable Emotion_Task_Acc]; gambling: mean of median RTs [variables Gambling_Task_Reward_Median_RT_Larger, Gambling_Task_Reward_Median_RT_Smaller, Gambling_Task_Punish_Median_RT_Larger, Gambling_Task_Punish_Median_RT_Smaller]; language: mean of story accuracy [variable Language_Task_Story_Acc], math accuracy [variable Language_Task_Math_Acc], story difficulty [variable Language_Task_Story_Avg_Difficulty_Level], and math difficulty [variable Language_Task_Math_Avg_Difficulty_Level]; relational: overall accuracy [variable Relational_Task_Acc]; social: mean accuracy [variables Social_Task_Random_Perc_Random and Social_Task_TOM_Perc_TOM]; WM: overall accuracy [variable WM_Task_Acc]) with gF for all participants in our sample who had performance data for that task (number of participants missing data: language, emotion, relational, WM: 0; social: 1; gambling: 20). We repeated this analysis 10 times for each task, using different data splits each time. This yielded 10 estimates of term predictive utility, and 10 estimates of gF relatedness for each task; mean and standard deviation of each are plotted in Fig. 2b and related via correlation. In the five tasks with accuracy data, very few (<=10) participants in our sample performed with below-chance accuracy in each task; excluding these subjects from these analyses did not change correlation results.

### Evaluating and visualizing contributions to a predictive model

Given the improved or comparable performance of the combined models relative to the individual-term models and the opportunity to interrogate relative term contributions to these combined models, we performed several analyses to evaluate the contributions of individual features, networks, and terms to the combined model for a given task. First, to ensure that only reliably predictive features were analyzed, a given feature was required to have been selected on 100% of all feature selections (10 folds * 90 iterations = 900 feature selections, for main analyses). The contribution of each selected feature was calculated as its mean ridge regression coefficients (b) across all 900 analyses multiplied by the standard deviation of its PPI beta across subjects. That is, the contribution ***c*** for feature *i* was defined as

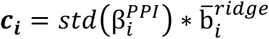

For all reliably selected features from a given term *t* (e.g., ciFC), the absolute values of these contributions were summed and converted to fractional contributions,

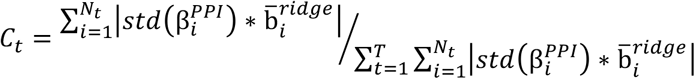

where *N_t_* represents the number of features from the given term *t* (with features that were not selected on 100% of feature selections set to 0) and *T* represents the number of terms. This procedure was repeated for each task to yield four contribution fractions per task (one per term). These contribution fractions were in turn used to drop terms in order of ascending and descending contributions, after which prediction was repeated to determine the impact of term contribution on model performance (Supplementary Fig. 2). Contributions across terms were compared using rank correlation of all features’ absolute contributions (ciFC and cdFC); of all nodes’ absolute contributions (cdFC contribution degree 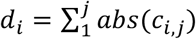, with *d_i_* the degree for node *i*, and *c_i,j_* the cdFC model contribution of the edge connecting nodes *i* and *j*) with absolute task activation contributions; and of absolute task activation contributions with absolute task activation, itself.

Contributions were visualized at several levels of analysis; results are displayed for the language task in Fig. 3 and for all tasks in Supplementary Fig. 4. First, signed contributions of each feature were visualized in circle plots (Fig. 3d-e, and Supplementary Fig. 4b, d). In these plots, nodes are grouped by canonical network (see *Functional parcellation and network definition*), and reliably selected edges are represented by lines between these nodes. The color family of each edge indicates the sign of its predictive contribution (red for positive, blue for negative), and the shade and thickness of the line represent the magnitude of its predictive contribution (darker and thicker indicate greater contribution). Fig. 3d and Supplementary Fig. 4b circle plots illustrate the predictive contributions of ciFC features for each task. Fig. 3e and Supplementary Fig. 4d circle plots illustrate the predictive contributions of context-dependent features with predictive utility. That is, lines again represent edges (here, cdFC), with the addition of node-level information. Nodes, displayed as circles on each circle plot track, are colored by their mean activation difference for the given contrast (e.g., 2-back – 0-back), and their distance from the x axis indicates their signed predictive contribution. To avoid any bias in task activation beta estimates from the inclusion of additional predictors in the PPI analysis (but see *Investigating potential confounds* and Supplementary Table 10 for evidence that PPI task activation betas closely follow independently estimated task effect sizes), activation was calculated for this analysis and for the inter-subject consistency/utility regression (see *Modeling the relationship between inter-subject consistency and predictive utility)* as follows. As in work by Salehi and colleagues^33^, we used individual-level, volume-based task contrast of parameter estimate (COPE) files generated and described previously^56^ using FSL FEAT’s FLAME (FMRIB’s Local Analysis of Mixed Effects)^62^ to calculate effect size for each voxel. One-sample *t*-statistics were calculated at each voxel and converted to Cohen’s *d* coefficients as 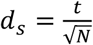, where *d_s_* is the sample *d* coefficient, *t* is the *t*-test statistic, and *N* is the sample size^63^. We then applied the initial 268-parcel group-level parcellation^28^ to these voxel-level maps to calculate a mean task effect size per parcel for each of the tasks.

Second, predictive contributions were visualized at the network level, using ten canonical networks (see *Functional parcellation and network definition;* Figs. 3a-c, 4a-b, 5b, and Supplementary Fig. 4a, c, e, f, and Supplementary Figs. 6-7). Given the potentially divergent interpretations of the predictive contribution of a positive and a negative edge or node (e.g., a positive contribution for a negative edge suggests that it is less negative [weaker] in those with higher fluid intelligence, while the same contribution for a positive edge suggests that it is more positive [stronger] in those with higher fluid intelligence), edges were first divided by their mean ciFC sign (i.e., edges that, at baseline, are on average positive across all subjects, and edges that, at baseline, are on average negative across all subjects; Fig. 3a-c, Supplementary Fig. 4a, c, f, and Supplementary Fig. 6), and nodes by their mean task activation sign (Fig. 3f, Supplementary Fig. 4e). For each group, the signed contributions (except for “diff” [Fig. 3c and Supplementary Fig. 4f], in which absolute contributions were used) of selected edges from each network pair were summed, and this sum was normalized by the total number of edges between the given networks to account for differences in network size, yielding the mean contribution for an edge in the given network pair. Finally, to increase interpretability of the scale for these network-level contributions, each network pair’s contribution value was normalized by the summed absolute contributions of all network pairs for that term in both positive and negative edge groups. This analysis was repeated, with minor modifications, for task activation for each network, rather than network pair. That is, activation predictive utility was summed for all nodes in each network, and this value was normalized by the number of nodes in that network; resulting network contributions were then normalized by the summed absolute contributions of all networks for that task in both positive and negative node groups. To explore any differences in the spatial distribution of high-consistency predictive edges and low-consistency predictive edges, this analysis was repeated after splitting the reliably selected edges not by mean ciFC sign, but rather by the median absolute inter-subject consistency (see *Inter-subject psychophysiological interaction analysis*) for these selected edges for the given term and task.

Finally, predictive contributions were visualized at the node level. Specifically, the difference between ciFC and cdFC predictive feature localization was further explored: for each task and term (ciFC and cdFC), absolute node degree was calculated (i.e., sum of absolute contributions for all edges incident to the given node) and normalized by total (i.e., summed) degree for that task/term to yield a fractional contribution. cdFC normalized degree was subtracted from ciFC normalized degree, yielding one 268×1 node difference vector per task; these vectors were averaged across tasks, rounded to the nearest integer, and scaled linearly for visualization (Fig. 3c). Similarly, for each task, absolute activation utility for each node was normalized by summed absolute activation utility for all nodes in the given task, and these node-level values were rounded to the nearest integer and linearly scaled for visualization (Fig. 3f).

### Inter-subject psychophysiological interaction analysis

Inter-subject FC isolates stimulus-induced inter-regional covariance from intrinsic and noise signals^26^; the diagonal of this FC matrix, termed inter-subject correlation (ISC), captures stimulus-induced intra-regional consistency of the BOLD signal^42^. To evaluate inter-subject consistency separately for ciFC, task activation, and cdFC, the intra-subject PPI analysis was repeated with one modification: the predictor node’s time course was averaged across the subset of all subjects that did not include the target subject (and that experienced the same task stimulus order; range across tasks = 608-703 subjects), and this process was repeated iteratively with each subject serving once as the target subject, yielding, again, one asymmetric matrix per term for each subject. That is,

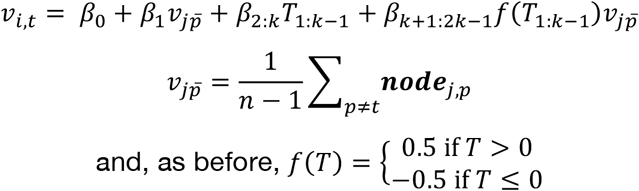

where *t* is the target subject, *p* are the non-target subjects, and *node_j,p_* represents the time course of the *j*^th^ node for subject *p* (Fig. 6). These matrices were averaged across all subjects, and, as before, contrasts were calculated, LR/RL contrast matrices averaged, and resulting beta contrast matrices symmetrized, yielding one matrix per term. Inter-subject consistency of task activation was defined as the main diagonal of the inter-subject interaction matrix. This was repeated for each of the five tasks for which task timing was meaningfully synchronized across subjects: gambling, emotion, WM, social, and relational. All subsequently described analyses using inter-subject consistency results were limited to these five tasks.

**Figure 6.**
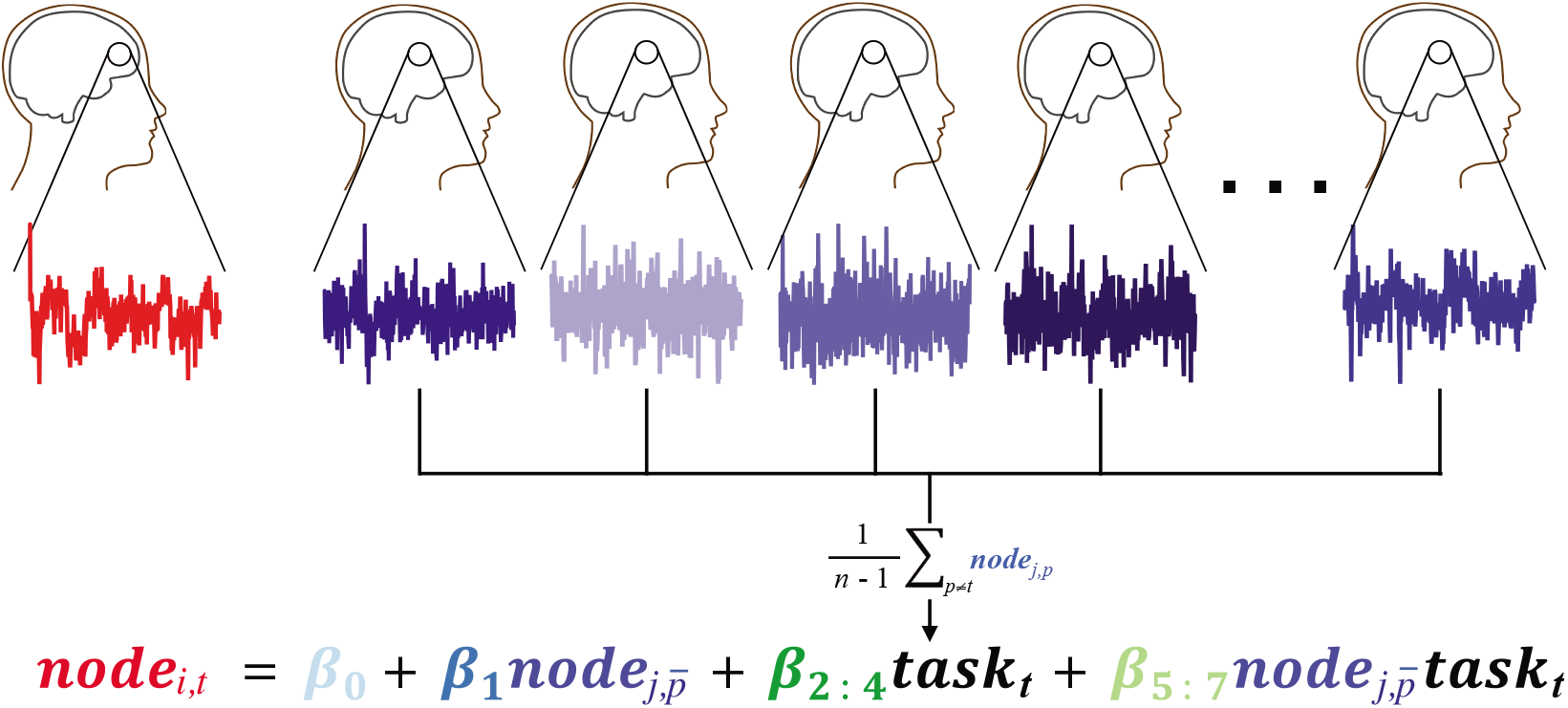
Schematic depiction of the inter-subject PPI analysis pipeline. The target node *i* time course is taken from subject *t*, and the predictor node *j* time course is averaged across all remaining subjects *p*.

To explore the spatial distribution of ciFC and cdFC inter-subject consistency, the mean consistency of an edge in each canonical network pair was visualized (Fig. 4a-b), using the same approach as in Figs. 3 and 5 (see *Evaluating and visualizing contributions to a predictive model*), again after dividing these edges into those with mean positive ciFC and those with mean negative ciFC (Supplementary Fig. 6c-d). We note that self-connections (e.g., node 1 – node 1) are constant in intra-subject analyses and thus non-contributory to predictive models. In inter-subject analyses, however, these self-connections correspond to inter-subject correlation (ISC)^42^, or the similarity in a given node’s time course across individuals. These connections are thus neither positive nor negative in the intra-subject analyses, but are included in Fig. 4a-b, and context-dependent ISC is interpreted as consistency of task activation across individuals in subsequent analyses.

### Modeling the relationship between inter-subject consistency and predictive utility

Given the substantial inter-subject consistency of ciFC, we sought to investigate a potential relationship between ciFC predictive utility and feature consistency across individuals. To do so, we designed a regression-based analysis in which the combined model predictive utility (i.e., absolute contribution) of each reliably selected ciFC feature for the given task was used as the outcome variable, and the inter-subject consistency of each corresponding ciFC feature (i.e., absolute inter-subject PPI beta) for the corresponding task was used as a predictor. We suspected that this relationship may interact with functional and anatomical relationships; to this end, we built five separate models, each with a different functional or anatomical variable explicitly modeled as a predictor: edge location within or between canonical networks (Supplementary Fig. 8c), edge location within or across hemispheres (Supplementary Fig. 8b), edge length (as measured by Euclidean distance between the nodes incident to that edge; Supplementary Fig. 8a), edge reliability (calculated in HCP resting-state data by Noble and colleagues^32^; Supplementary Fig. 8d), and task activation (i.e., mean absolute task effect size [see *Evaluating and visualizing contributions to a predictive model*] of the nodes incident to the given edge; Fig. 5c). All non-dummy predictors were mean-centered (i.e., *z*-scored). Both main effects and interaction terms of models with significant full-model *P* values are presented; betas of non-significant models were set to 0. Due to the limited number of measurements (i.e., five betas per model term, one per task) output by this analysis, results are discussed qualitatively, with the caveat that future work to replicate these findings with more tasks will permit more rigorous statistical testing of them (see *Discussion*).

### Investigating potential confounds

To evaluate the potential impact of collinearity on PPI beta estimates, intra- and inter-subject PPI analyses were repeated with two partial models for each task: one without the ciFC term and one without the cdFC term. Partial-model and full-model betas for each term were highly correlated, suggesting a minimal impact of collinearity on beta estimates (Supplementary Table 8 and Supplementary Table 9).

To ensure that ciFC and task activation beta estimates are comparable to standard measures of FC and task activation, respectively, ciFC betas were correlated with FC calculated using Fisher-transformed Pearson correlations across the entire node time courses (“standard FC”), mean task activation betas were correlated with group-level task effect size measures calculated for each node (see *Evaluating and visualizing contributions to a predictive model)*, and HCP-released, individual-level GLM results were correlated with PPI task activation betas and averaged across subjects, for all subjects in the main sample for whom individual-level GLM results were available (*n* = 322; all results in Supplementary Table 10). These individual-level GLM results were also used for prediction, to ensure that results were comparable to prediction performance using PPI-based task activation (Supplementary Table 10). For completeness, cdFC beta estimates were also correlated with standard FC. As predicted, results (Supplementary Table 10) demonstrate that ciFC is strongly correlated with standard FC and task activation betas are comparable to HCP-released, GLM-based task effect size (both at the group and individual levels), but that cdFC is not significantly correlated with standard FC, validating the interpretation of PPI betas.

To further ensure that methodological choices did not affect main results, we repeated main analyses with mean-centered PPI regressors (that is, after *z*-scoring each PPI predictor immediately prior to the regression step), and non-centered task regressors (that is, condition on = 1, condition off = 0). Prediction results were largely unchanged (Supplementary Fig. 1 and Supplementary Table 11), though the intercept term, by definition, failed to predict gF in the mean-centered case (numerical error will yield intercept values that are close, but not equal, to zero, but this fluctuation around zero should not—and did not—predict gF). While at first surprising that prediction results are comparable using zero- and mean-centered approaches, it follows from the similarity of task timing across subjects and from our choice to *z*-score node timecourses. That is, mean centering PPI regressors causes a linear scaling of resulting betas that is comparable across subjects, since 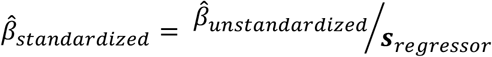, and the standard deviation of task timing and of the interaction will be nearly identical across subjects. This linear shift in PPI betas will change their interpretation, but this change is not germane to the present work, as PPI betas themselves are not interpreted. However, with the exception of the intercept term, PPI betas’ predictive utility will be unchanged, since inter-subject relationships of PPI betas are relatively unchanged. Further, predictive utility estimates will be essentially unchanged, as ridge coefficients scale with PPI beta variance. The intercept will, of course, approach zero in the mean-centered case, but in the zero-centered case will reflect these linear shifts, scaled by subject-specific PPI activation and cdFC betas (Supplementary Fig. 1). If these betas meaningfully vary across subjects, then the intercept terms may predict individuals’ phenotypes, as was found to be true in these analyses (Fig. 2). Given the predictive utility of this “overall task effect” term (i.e., intercept), the unchanged prediction results for other terms, and the decreased collinearity among predictors after zero centering, we present zero-centered results in the main text, and mean-centered results in the supplementary materials.

Next, to ensure that sample particularities and QC issues did not contribute to results, we repeated main analyses after excluding all subjects who experienced HCP QC issue C or recon version r177 (*n* = 531). Prediction performance was comparable to main results (Supplementary Table 12).

Finally, while standard approaches were taken to mitigate the effects of motion on fMRI data, we sought to more thoroughly explore any relationship of motion to task timing by correlating, for each subject, frame-to-frame displacement (HCP Movement_RelativeRMS.txt) for each task and phase encoding direction with the corresponding task timing regressors. Results (Supplementary Table 6) demonstrate no consistent relationship. We also correlated mean RMS relative motion for each subject and task (averaged over phase encoding runs) with observed gF (Supplementary Table 2) and predicted gF (averaged over 90 iterations for each task/term; Supplementary Table 3). Given several modest correlations, we repeated the prediction analyses using partial correlation-based feature selection with mean RMS relative motion (calculated for each subject and task) as a covariate. As an even more conservative motion control analysis, we also repeated the main analysis after regressing mean RMS relative motion out of gF and FC within the cross-validation loop. Regression coefficients were estimated for the training subjects and applied to the test subjects in the 10-fold analysis (to avoid the use of potentially unstable coefficient estimates in the smaller test sample), but were estimated separately for training and test subjects in the split-half analysis. Model performance and feature weights were relatively unchanged; the former is presented in Supplementary Table 4, and the correlations of feature weights from main analyses and partial correlation-based analyses are presented in Supplementary Table 5. Correlations of feature weights from main analyses and residual-based analyses were comparable. Finally, we repeated main prediction analyses using the half of the sample with the lowest grand mean RMS relative motion (< 0.072mm across tasks; *n* = 351). Results are presented in Supplementary Table 4.

### Data availability

The HCP data that support the findings of this study are publicly available on the ConnectomeDB database (https://db.humanconnectome.org). Data used to generate the 268-node parcellation and to define the canonical networks can be found at http://fcon_1000.projects.nitrc.org/indi/retro/yale_hires.html.

### Code availability

MATLAB code to run the ridge regression-based CPM analysis can be found at https://github.com/YaleMRRC/CPM. MATLAB code to run additional core analyses (PPI analyses, basic visualization, synchrony vs. predictiveness analyses, and family-based cross-validation) can be found at https://github.com/abigailsgreene/taskFC. BioImage Suite tools used for analysis and visualization can be accessed at www.bisweb.yale.edu. MATLAB and R scripts to perform additional post-hoc analyses and visualizations are available from the authors upon request.

## Acknowledgments

Data were provided by the Human Connectome Project, WU-Minn Consortium (Principal Investigators: David Van Essen and Kamil Ugurbil; 1U54MH091657) funded by the 16 NIH Institutes and Centers that support the NIH Blueprint for Neuroscience Research; and by the McDonnell Center for Systems Neuroscience at Washington University. This work was also supported by T32GM007205 (A.S.G.). The authors would like to thank Nick Turk-Browne, Daeyeol Lee, and Daniel Barson for helpful discussions and suggestions.

## Author contributions

R.T.C., D.S., and A.S.G. designed the study. S.N. developed the scripts used to preprocess the HCP data, and A.S.G. preprocessed the data with guidance from S.N. S.G. developed and validated the ridge regression-based predictive modeling pipeline. A.S.G. developed and validated the PPI and visualization code and performed all analyses, with the exception of task effect size calculation (designed and overseen by S.N.). D.S. and R.T.C. supported result interpretation. A.S.G. wrote the manuscript, with comments from all authors.

