## Supplementary Figures for "How tasks change whole-brain functional organization to reveal brain-phenotype relationships"

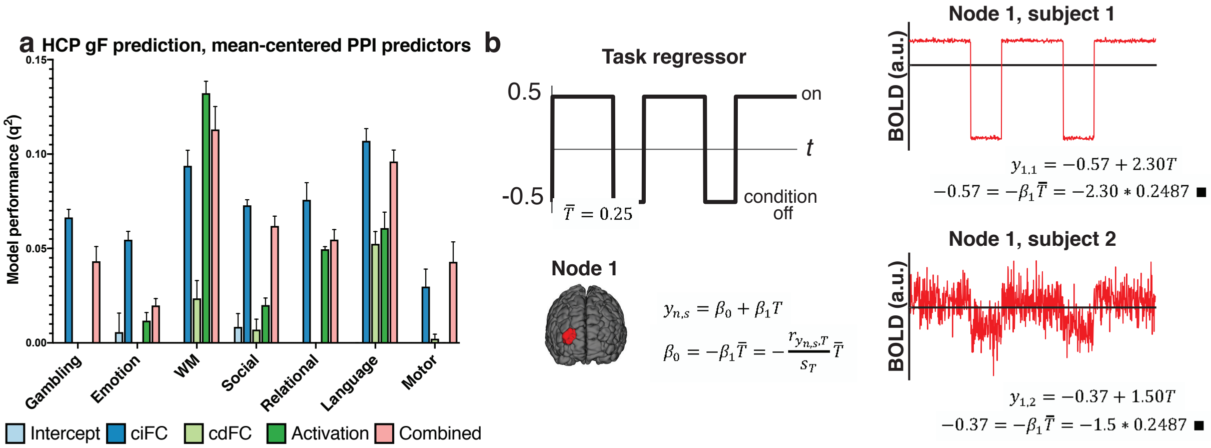


**Supplementary Figure 1.** PPI intercept reflects overall degree of task effect on brain activity patterns, and has predictive utility. (a) Eliminating PPI model intercepts by mean centering all regressors (rather than zero centering task regressors and mean centering node time courses, as in main analyses) has little effect on prediction performance (mean and s.d., indicated by error bars) of ciFC, cdFC, activation, or combined models (c.f., Fig. 2a, Supplementary Fig. 3) because removal of a constant term (that is similar across subjects) from regressors scales beta estimates in a linear fashion, and thus does not affect their phenotypically relevant information content. However, the intercept reflects this constant term, weighted by activation and interaction betas, which may vary across subjects, reflecting inter-individual differences in overall degree of task effect on node activity and FC; should this inter-individual variance be related to gF, PPI intercept terms would be expected to successfully predict gF, as was found to be the case for 5/7 tasks (Fig. 2). (b) Schematic illustration of this interpretation of the intercept term. Here, task timing is represented as a zero-centered boxcar, yielding a mean task regressor value of 0.25 (because more time is spent in the "on" condition than in the "off" condition). Example node 1 time courses for two subjects are depicted in the rightmost panel; in subject 1, node 1 is strongly activated by the task, while in subject 2, node 1 is more weakly activated by the task, as reflected in the $\beta_1$ estimates for these subjects. This difference, in turn, determines the intercept values for each subject. This simple case excludes the PPI interaction term, but the same logic would hold for this term, rendering the intercept a reflection of the overall degree to which the task affects activation and connectivity of the target node, a value which may vary meaningfully across individuals. T, task regressor; y, node time course of activity; n, node; s, subject; *r*, Pearson correlation; *s*, standard deviation.

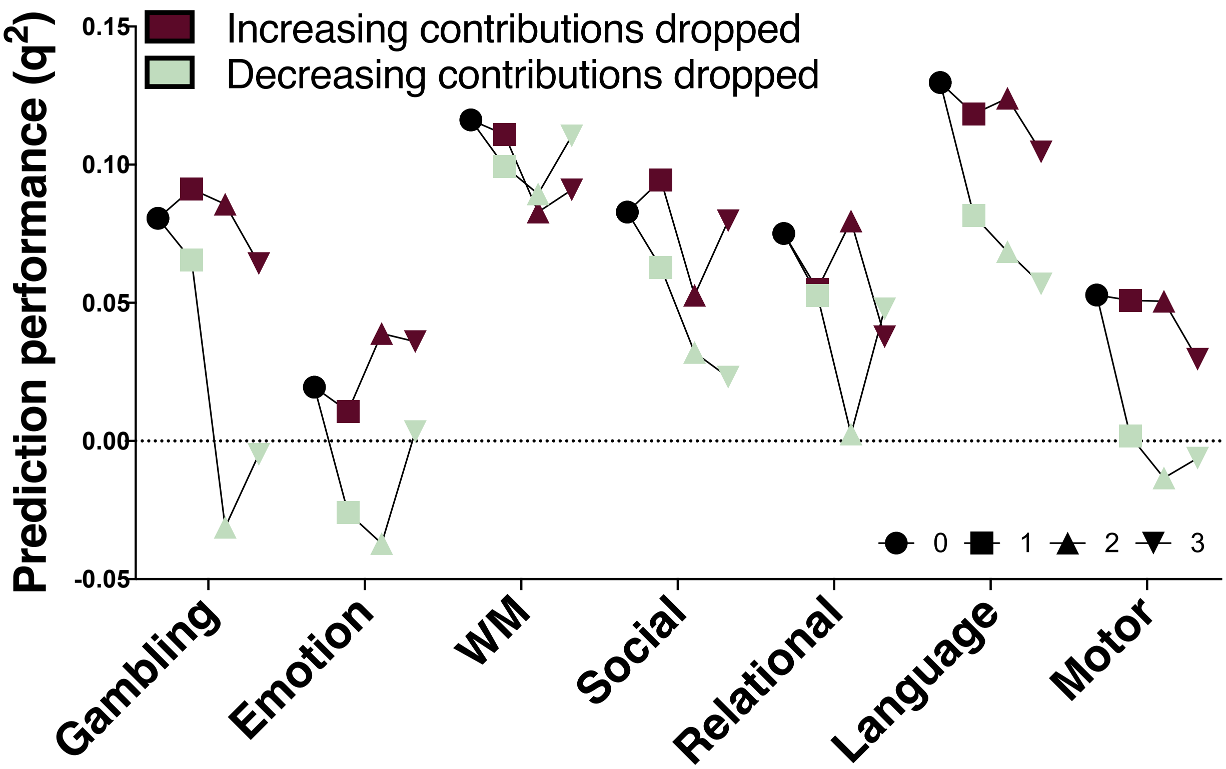


**Supplementary Figure 2.** Combined model prediction performance with increasing numbers of individual terms dropped from the model in order of increasing (red) and decreasing (green) contributions. Numbers indicate number of dropped terms.

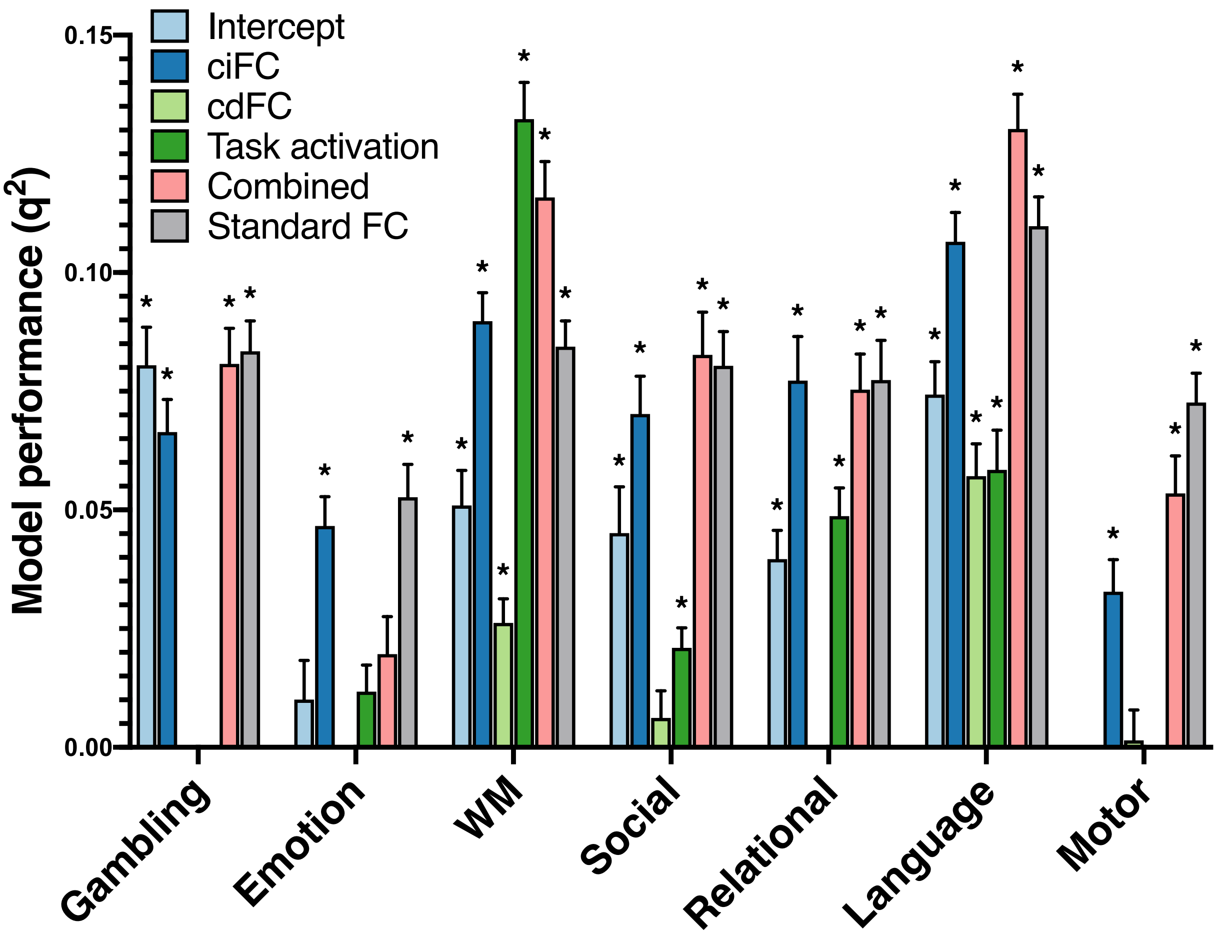


**Supplementary Figure 3.** Summary (mean and s.d., indicated by error bars) of model performance for all terms and tasks (*$P$ < 0.01, corrected, via permutation test). Models with negative mean $q^2$ were set to 0. “Standard FC”: prediction performance using FC calculated as Fisher-transformed Pearson correlations, without task modeling (see Methods).

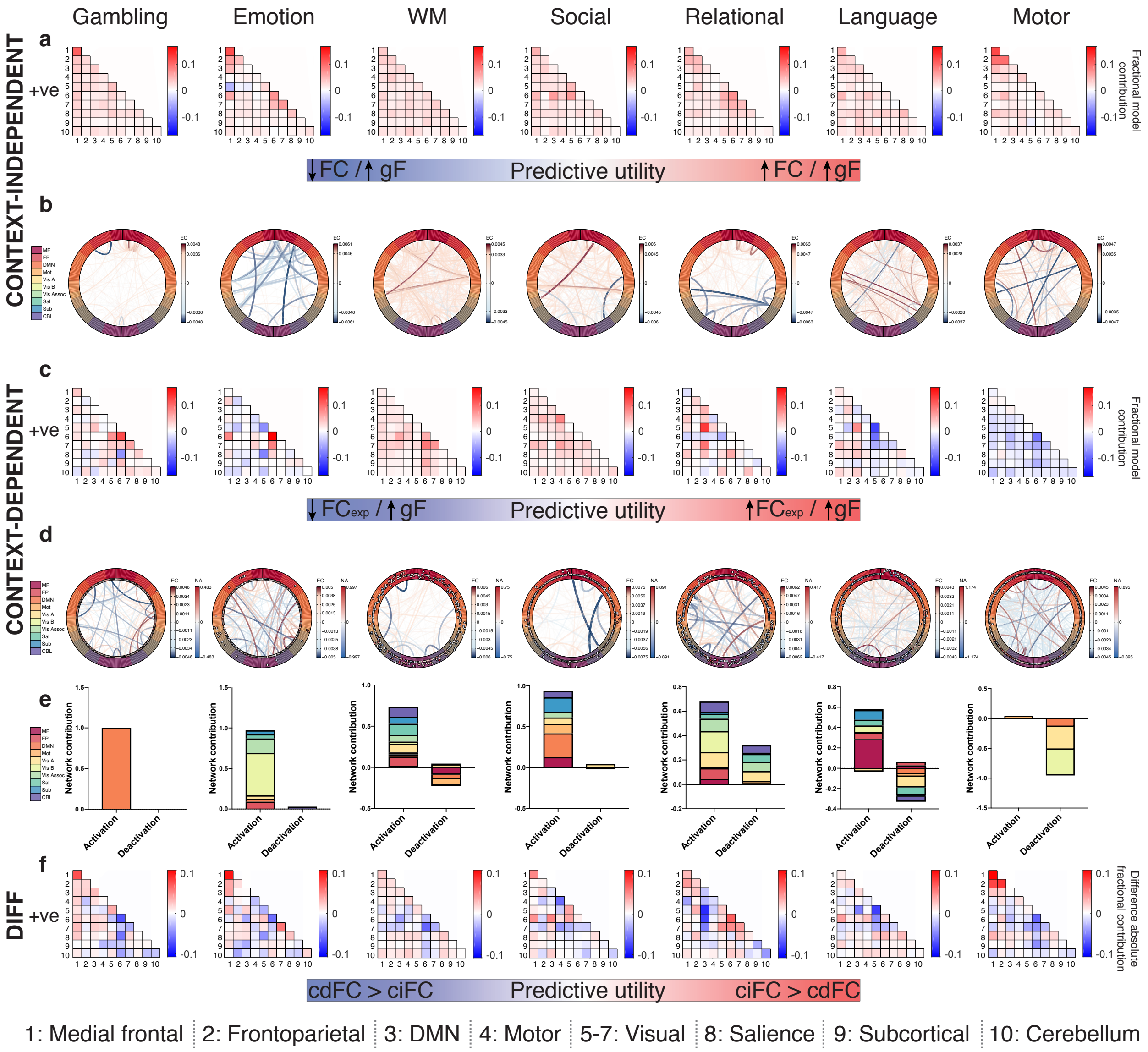


**Supplementary Figure 4. Context-independent FC, context-dependent FC, and activation predictive features are distributed and distinct.** (a) Visualization of predictive ciFC features by network for each task. Red = positive mean ridge coefficients, blue = negative mean ridge coefficients, shade = relative model contribution. In this and all subsequent figures: "+ve" indicates that results reflect only contributions of edges with mean positive ciFC. 1-10 = network assignment. (b) Visualization of individual predictive ciFC features, with each consistently selected edge represented as a line; line color and thickness scale with predictive model contribution. In this and all subsequent figures, MF = medial frontal, FP = frontoparietal, DMN = default mode network, Mot = motor, Vis A = visual A, Vis B = visual B, Vis Assoc = visual association, Sal = salience, Sub = subcortical, CBL = cerebellum. EC, edge contribution. (c) Visualization of predictive cdFC features by network for each task. $FC_{exp}$, FC during the experimental condition (i.e., condition of interest). Red = positive mean ridge coefficients, blue = negative mean ridge coefficients, shade = relative model contribution. 1-10 = network assignment. (d) Visualization of individual predictive cdFC (lines) and activation (outer track circles) features; line color and thickness scale with cdFC feature predictive utility, and circles represent the corresponding nodes, with their color indicating mean activation (red = positive, blue = negative) and distance from the x axis indicating their model contribution. EC, edge contribution; NA, node activation. (e) Visualization of fractional mean network contributions for predictive nodes with mean positive PPI activation betas (“activation”) and for nodes with mean negative PPI activation betas (“deactivation”). Note that two tasks (gambling and motor) had fewer than 20 consistently predictive nodes, rendering network visualizations potentially difficult to interpret. (f) Visualization of the difference between absolute contribution-based, network-level ciFC model contributions and absolute contribution-based, network-level cdFC model contributions.

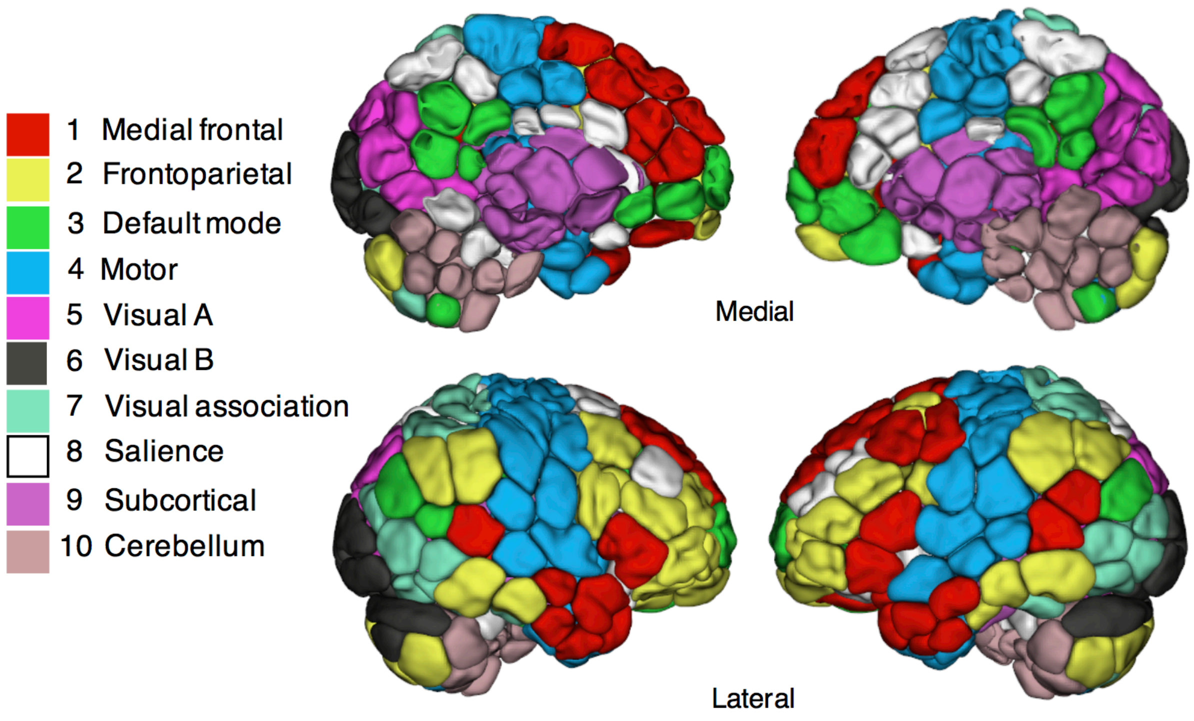


**Supplementary Fig. 5.** Ten canonical networks used for network analyses (see Methods for derivation). Figure adapted from (Greene, Gao, Scheinost, & Constable, 2018).

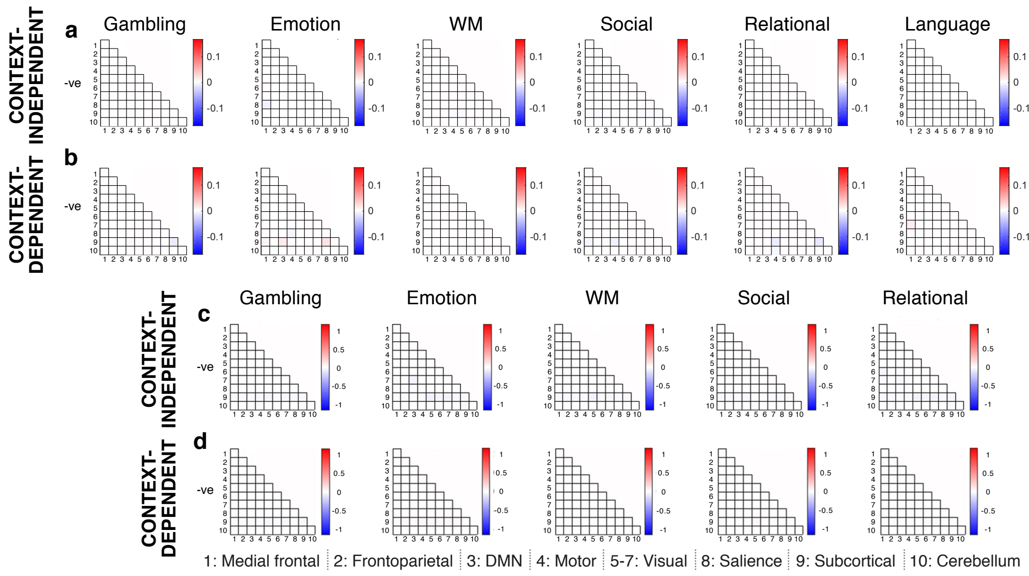


**Supplementary Figure 6.** Predictive model contributions by network of features with mean negative ciFC (a,b) and inter-subject consistency of features with mean negative ciFC (c,d), summarized by canonical networks (Fig. S5). Note that the number of selected edges with mean negative ciFC (a,b) ranged from 0-42 across tasks.

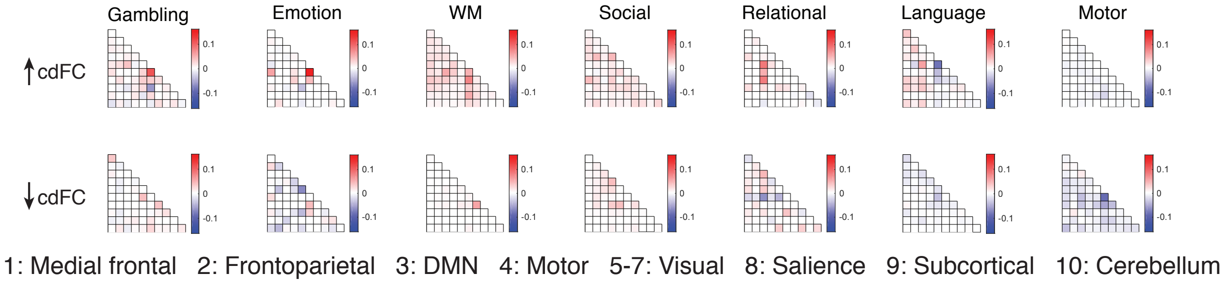


**Supplementary Figure 7.** Further separating cdFC model contributions (for features with positive mean ciFC) by positive (top row) and negative (bottom row) cdFC offers additional insight into predictive task-induced changes in FC.

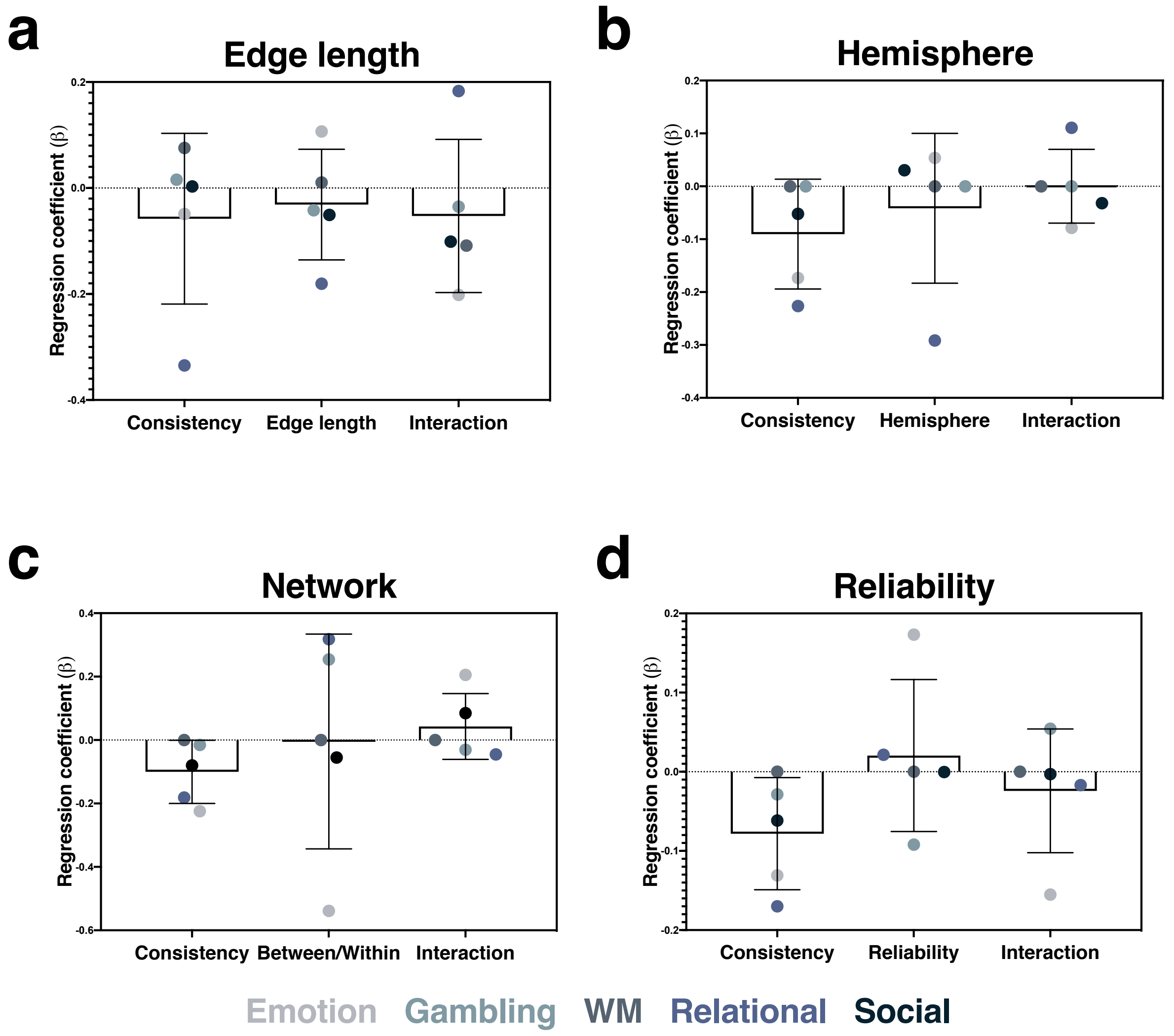


**Supplementary Figure 8.** Relationships between inter-subject consistency, edge length (a), brain hemisphere (b), network membership (c), reliability (d), and predictive utility. Bar height reflects mean coefficient; error bars indicate s.d. of coefficients across tasks.

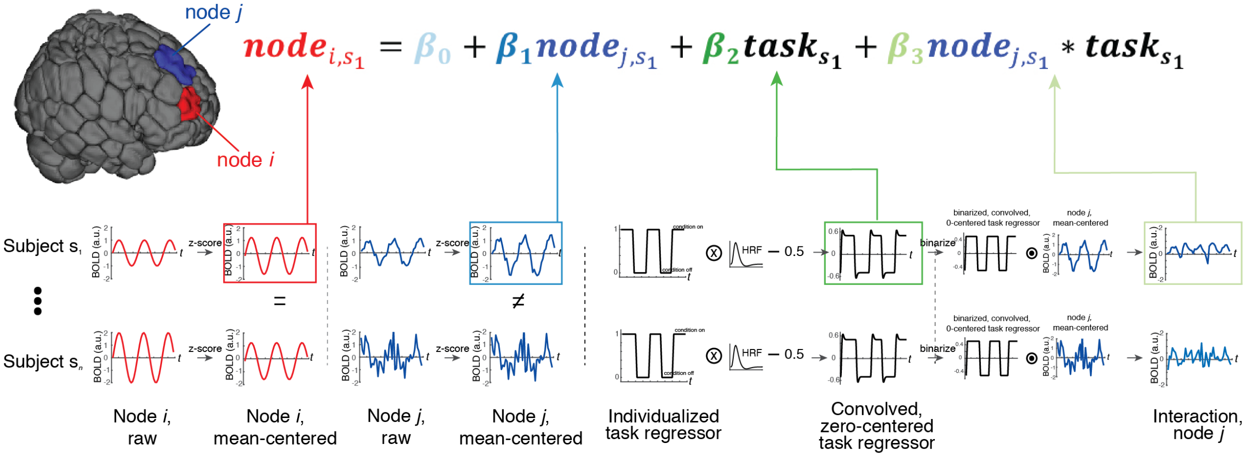


**Supplementary Figure 9.** Schematic depiction of the PPI analysis pipeline.

| | Prediction performance, no GSR | Prediction performance, with GSR |
|---|---|---|
| **Gambling** | 0.0834 (0.006) | 0.1084 (0.007) |
| **Emotion** | 0.0527 (0.007) | 0.0848 (0.006) |
| **WM** | 0.0844 (0.005) | 0.1247 (0.007) |
| **Social** | 0.0804 (0.007) | 0.0973 (0.007) |
| **Relational** | 0.0774 (0.008) | 0.1237 (0.008) |
| **Language** | 0.1098 (0.006) | 0.1087 (0.006) |
| **Motor** | 0.0726 (0.006) | 0.0886 (0.007) |

**Supplementary Table 1.** Prediction performance of rCPM (90 iterations, *P* threshold = 0.1) performed on "standard FC," with and without global signal regression (GSR). Results presented as mean $q^2$ (s.d. $q^2$). We note that model performance without GSR is overall lower than previously reported (Greene et al., 2018), a finding that holds when standard FC was passed through the same preprocessing and prediction pipelines (Supplementary Fig. 3). However, the addition of GSR to the FC matrix preprocessing pipeline substantially improved prediction performance, rendering it comparable to previous results. While GSR was not appropriate for main analyses due to potential task-related fluctuations of the global signal and corresponding, interpretable effects on the task activation signal (e.g., due to fluctuating arousal or vigilance [Liu et al., 2017]), and while these analyses depend on relative, rather than absolute, prediction performance, this finding supports the utility of GSR for FC-based prediction (Greene et al., 2018; Li et al., 2019).

| | Motion/observed gF correlation |
|---|---|
| **Emotion** | -0.14* |
| **Gambling** | -0.06 |
| **Language** | -0.09 |
| **Motor** | -0.10 |
| **Relational** | -0.15* |
| **Social** | -0.12* |
| **WM** | -0.08 |

**Supplementary Table 2.** Pearson correlation between mean RMS motion (averaged between LR and RL runs for each subject) and observed gF. *indicates significant at $P<0.05$, Bonferroni corrected.

| | Intercept | ciFC | cdFC | Activation | Combined |
|---|---|---|---|---|---|
| **Emotion** | -0.0076 | -0.1702*+ | -0.0211 | -0.0694 | -0.1129 |
| **Gambling** | 0.0057+ | -0.0536+ | -0.0041 | 0.0576 | -0.0274+ |
| **Language** | -0.1614*+ | -0.1181+ | -0.1482*+ | -0.0752+ | -0.1590*+ |
| **Motor** | -0.0296 | -0.2176*+ | 0.0199 | -0.0599 | -0.1333*+ |
| **Relational** | -0.1015+ | -0.1158+ | -0.0063 | -0.1228*+ | -0.1012+ |
| **Social** | -0.0524+ | -0.0979+ | -0.0158 | -0.1203*+ | -0.0575+ |
| **WM** | -0.0666+ | -0.1002+ | -0.0772+ | -0.1451*+ | -0.0798+ |

**Supplementary Table 3.** Pearson correlation between mean RMS motion (averaged between LR and RL runs for each subject) and predicted gF (averaged across 90 iterations; see main text) for all tasks and terms. *indicates significant at $P<0.05$, Bonferroni corrected; +indicates significant corresponding predictive model (see Fig. 2).

| | | Intercept | ciFC | cdFC | Activation | Combined |
|---|---|---|---|---|---|---|
| **Emotion** | *Parcorr (10/1)* | 0.0018 | 0.0593 | -0.0351 | 0.0055 | 0.0203 |
| | *Residualized (10/1)* | 0.0146 | 0.0531 | -0.0446 | 0.0154 | 0.0222 |
| | *Original (10/90)* | 0.0100 | 0.0466 | -0.0394 | 0.0117 | 0.0196 |
| | *Residualized (2/90)* | -0.0082 | 0.0152 | -0.0420 | 0.0013 | -0.0096 |
| | *Lowest motion (10/5)* | 0.0366 | 0.0347 | -0.0543 | -0.0134 | 0.0255 |
| **Gambling** | *Parcorr (10/1)* | 0.0795 | 0.0686 | -0.0209 | -0.0110 | 0.0851 |
| | *Residualized (10/1)* | 0.0832 | 0.0666 | -0.0275 | -0.0036 | 0.0797 |
| | *Original (10/90)* | 0.0804 | 0.0664 | -0.0216 | -0.0065 | 0.0807 |
| | *Residualized (2/90)* | 0.0453 | 0.0368 | -0.0344 | -0.0101 | 0.0462 |
| | *Lowest motion (10/5)* | 0.0335 | 0.0056 | -0.0386 | -0.0134 | 0.0241 |
| **Language** | *Parcorr (10/1)* | 0.0678 | 0.1167 | 0.0568 | 0.0517 | 0.1218 |
| | *Residualized (10/1)* | 0.0703 | 0.0945 | 0.0535 | 0.0557 | 0.1243 |
| | *Original (10/90)* | 0.0742 | 0.1065 | 0.0571 | 0.0585 | 0.1302 |
| | *Residualized (2/90)* | 0.0448 | 0.0681 | 0.0296 | 0.0567 | 0.0926 |
| | *Lowest motion (10/5)* | 0.0281 | 0.0486 | -0.0120 | 0.0330 | 0.0575 |
| **Motor** | *Parcorr (10/1)* | -0.0289 | 0.0280 | -0.0061 | -8E-4 | 0.0402 |
| | *Residualized (10/1)* | -0.0146 | 0.0360 | 0.0021 | -0.0019 | 0.0468 |
| | *Original (10/90)* | -0.0143 | 0.0328 | 0.0015 | -0.0018 | 0.0535 |
| | *Residualized (2/90)* | -0.0258 | 0.0147 | -0.0187 | -0.0073 | 0.0292 |
| | *Lowest motion (10/5)* | -0.0237 | -0.0063 | 0.0294 | -0.0027 | 0.0342 |
| **Relational** | *Parcorr (10/1)* | 0.0485 | 0.0742 | -0.0162 | 0.0510 | 0.0829 |
| | *Residualized (10/1)* | 0.0469 | 0.0797 | -0.0096 | 0.0434 | 0.0829 |
| | *Original (10/90)* | 0.0396 | 0.0772 | -0.0168 | 0.0487 | 0.0754 |
| | *Residualized (2/90)* | 0.0202 | 0.0288 | -0.0330 | 0.0414 | 0.0359 |
| | *Lowest motion (10/5)* | 0.0055 | 0.0389 | -0.0250 | 0.0174 | 0.0279 |
| **Social** | *Parcorr (10/1)* | 0.0587 | 0.0680 | 0.0118 | 0.0164 | 0.0888 |
| | *Residualized (10/1)* | 0.0421 | 0.0750 | 0.0065 | 0.0236 | 0.0917 |
| | *Original (10/90)* | 0.0451 | 0.0702 | 0.0062 | 0.0210 | 0.0827 |
| | *Residualized (2/90)* | 0.0085 | 0.0342 | -0.0048 | 0.0085 | 0.0445 |
| | *Lowest motion (10/5)* | -0.0068 | 0.0324 | 0.0080 | 0.0359 | 0.0358 |
| **WM** | *Parcorr (10/1)* | 0.0598 | 0.0911 | 0.0254 | 0.1373 | 0.1169 |
| | *Residualized (10/1)* | 0.0542 | 0.0895 | 0.0293 | 0.1258 | 0.1137 |
| | *Original (10/90)* | 0.0509 | 0.0898 | 0.0262 | 0.1323 | 0.1159 |
| | *Residualized (2/90)* | 0.0357 | 0.0566 | 0.0110 | 0.0934 | 0.0869 |
| | *Lowest motion (10/5)* | 0.0340 | 0.0436 | 0.0005 | 0.0592 | 0.0617 |

**Supplementary Table 4.** Comparison of main results ("Original") to performance of predictive models built using partial correlation-based feature selection ("Parcorr," i.e., controlling for motion), regression of motion values out of FC and gF ("Residualized"), and only the half of the sample (*n*=351) that moved the least (grand mean RMS relative motion < 0.072mm across tasks). In all cases, *P* threshold = 0.1. Numbers in parentheses indicate number of folds and iterations (e.g., "Parcorr (10/1)" indicates partial correlation-based feature selection with 10 folds and 1 iteration). For cases in which a single iteration of prediction was performed, results presented as $q^2$; for cases in which more than one iteration of prediction was performed, results presented as mean $q^2$.

| | Intercept | ciFC | cdFC | Activation | Combined |
|---|---|---|---|---|---|
| **Emotion** | 0.5914 | 0.5867 | 0.4335 | 0.7402 | 0.5712 |
| **Gambling** | 0.6529 | 0.7503 | 0.4157 | 0.5795 | 0.6987 |
| **Language** | 0.6840 | 0.7534 | 0.5925 | 0.7908 | 0.7150 |
| **Motor** | 0.4247 | 0.6702 | 0.4935 | 0.5752 | 0.5862 |
| **Relational** | 0.7519 | 0.6096 | 0.4090 | 0.9309 | 0.6675 |
| **Social** | 0.5265 | 0.7321 | 0.5745 | 0.7101 | 0.6681 |
| **WM** | 0.7568 | 0.7888 | 0.6728 | 0.9137 | 0.7480 |

**Supplementary Table 5.** Rank correlation between consistently selected features' rCPM betas for the main models and the partial correlation-based models.

| | Task timing/motion, LR correlation | Task timing/motion, RL correlation |
|---|---|---|
| **Emotion** | -0.0002 | 0.0030 |
| **Gambling** | -0.0167 | -0.0220 |
| **Language** | -0.0132 | -0.0124 |
| **Motor** | 0.0141 | 0.0065 |
| **Relational** | 0.0011 | -0.0048 |
| **Social** | -0.0292 | -0.0344 |
| **WM** | -0.0055 | -0.0015 |

**Supplementary Table 6.** Rank correlation coefficients between frame-to-frame relative RMS motion and task timing, averaged across task conditions and subjects.

| | | Intercept | ciFC | cdFC | Activation | Combined |
|---|---|---|---|---|---|---|
| **Gam** | $q^2$ | **0.0804** | **0.0664** | -0.0216 | -0.0065 | **0.0807** |
| | $r$ | **0.2887** | **0.2605** | -0.0048 | -0.1134 | **0.2858** |
| **Emo** | $q^2$ | 0.0100 | **0.0466** | -0.0394 | 0.0117 | 0.0196 |
| | $r$ | 0.1229 | **0.2208** | -0.0786 | 0.1177 | 0.1614 |
| **WM** | $q^2$ | **0.0509** | **0.0898** | **0.0262** | **0.1323** | **0.1159** |
| | $r$ | **0.2281** | **0.3007** | **0.1731** | **0.3765** | **0.3447** |
| **Soc** | $q^2$ | **0.0451** | **0.0702** | 0.0062 | **0.0210** | **0.0827** |
| | $r$ | **0.2256** | **0.2689** | 0.1232 | **0.1465** | **0.2969** |
| **Rel** | $q^2$ | **0.0396** | **0.0772** | -0.0168 | **0.0487** | **0.0754** |
| | $r$ | **0.2033** | **0.2800** | 0.0116 | **0.2215** | **0.2777** |
| **Lang** | $q^2$ | **0.0742** | **0.1065** | **0.0571** | **0.0585** | **0.1302** |
| | $r$ | **0.2736** | **0.3272** | **0.2397** | **0.2686** | **0.3611** |
| **Mot** | $q^2$ | -0.0143 | **0.0328** | 0.0015 | -0.0018 | **0.0535** |
| | $r$ | 0.0830 | **0.1881** | 0.1050 | 0.0380 | **0.2424** |

**Supplementary Table 7.** Patterns of prediction performance across terms and tasks are comparable using MSE-based ($q^2$) and correlation-based (*r;* predicted versus observed gF) performance metrics. Bolded values indicate models with significant $q^2$ via non-parametric permutation tests.

| | Intercept, no cdFC | Intercept, no ciFC | ciFC, no cdFC | Activation, no cdFC | Activation, no ciFC | cdFC, no ciFC |
|---|---|---|---|---|---|---|
| **Gambling** | 0.9910 | 0.9587 | 0.969 | 0.9941 | 0.9749 | 0.7439 |
| **Emotion** | 0.9892 | 0.9654 | 0.9918 | 0.9903 | 0.9569 | 0.9343 |
| **Language** | 0.9779 | 0.9337 | 0.9617 | 0.9877 | 0.9558 | 0.7679 |
| **Motor** | 0.9754 | 0.9961 | 0.5986 | 0.9751 | 0.9915 | 0.7329 |
| **Relational** | 0.9821 | 0.9438 | 0.9881 | 0.9878 | 0.9542 | 0.9290 |
| **Social** | 0.9868 | 0.9275 | 0.9661 | 0.9925 | 0.9570 | 0.7050 |
| **WM** | 0.9848 | 0.9228 | 0.9862 | 0.9891 | 0.9548 | 0.8920 |

**Supplementary Table 8.** Mean Pearson correlation coefficients between full-model PPI betas and partial-model PPI betas for a given term, computed within subject (separately for LR and RL runs) and averaged across subjects and then conditions (e.g. cue, 2-back, and 0-back terms for WM).

| | Intercept, no cdFC | Intercept, no ciFC | ciFC, no cdFC | Activation, no cdFC | Activation, no ciFC | cdFC, no ciFC |
|---|---|---|---|---|---|---|
| **Gambling** | 0.9830 | 0.7846 | 0.9368 | 0.9659 | 0.8441 | 0.9075 |
| **Emotion** | 0.9563 | 0.9045 | 0.9407 | 0.9477 | 0.8550 | 0.9370 |
| **Relational** | 0.9604 | 0.8388 | 0.9503 | 0.9428 | 0.8066 | 0.9655 |
| **Social** | 0.9749 | 0.7681 | 0.9308 | 0.9642 | 0.8190 | 0.8768 |
| **WM** | 0.9468 | 0.8164 | 0.9647 | 0.9515 | 0.8643 | 0.9684 |

**Supplementary Table 9.** Mean Pearson correlation coefficients between inter-subject PPI betas for full and partial models.

| | ciFC/ standard FC corr | cdFC/ standard FC corr | Act. PPI/GLM grp corr | Act. PPI/GLM IS corr | GLM-based model performance |
|---|---|---|---|---|---|
| **Emotion** | 0.973 | -0.0975 (0.25) | 0.9360 | 0.9196 | -0.0194 |
| **Gambling** | 0.9364 | -0.0049 (0.35) | 0.7798 | 0.6725 (0.0001) | -0.0146 |
| **Language** | 0.9512 | 0.0206 (0.39) | 0.9363 | 0.9369 | 0.0669 |
| **Motor** | 0.6499 | -0.0717(0.43) | 0.8897 | 0.8259 (0.0001) | -0.0206 |
| **Relational** | 0.9546 | -0.0123 (0.34) | 0.9457 | 0.7760 | 0.0652 |
| **Social** | 0.9278 | -0.0064 (0.42) | 0.92 | 0.8198 | 0.0110 |
| **WM** | 0.9554 | -0.0428 (0.35) | 0.9472 | 0.8448 (0.0016) | 0.0965 |

**Supplementary Table 10.** Comparison of PPI results to "standard" FC and activation results. Columns 1-2: Pearson correlations of context-independent and context-dependent FC for each task with standard FC from that task, computed within subject and averaged across subjects to yield mean *r*(*P*, Bonferroni corrected). *P* value not reported indicates *P* << 0.001. Column 3: Pearson correlations of task activation PPI betas (averaged across subjects) with independently estimated, group-level ("grp") HCP task effect sizes per node. All *P* << 0.001, Bonferroni corrected. Column 4: Pearson correlations of task activation PPI betas with independently estimated, individual-level HCP task effect sizes per node; intra-subject ("IS") correlations averaged across subjects and presented as mean *r(P*, Bonferroni corrected). *P* value not reported indicates *P* << 0.001. Act, activation; corr, correlation; grp, group. Column 5: performance of gF predictive models trained and tested with GLM-based activation, rather than PPI-based activation. Results reported as mean $q^2$ across 5 iterations of 10-fold cross-validation.

| | | Intercept | ciFC | cdFC | Activation | Combined |
|---|---|---|---|---|---|---|
| **Emo** | *Mean-centered* | 0.0057 | 0.0546 | -0.0395 | 0.0118 | 0.0198 |
| | *Non-centered* | 0.0328 | 0.0310 | -0.0398 | 0.0117 | 0.0173 |
| **Gam** | *Mean-centered* | -0.0054 | 0.0664 | -0.0221 | -0.0056 | 0.0432 |
| | *Non-centered* | 0.0868 | 0.0249 | -0.0224 | -0.0055 | 0.0753 |
| **Lang** | *Mean-centered* | -0.0029 | 0.1070 | 0.0525 | 0.0608 | 0.0960 |
| | *Non-centered* | 0.1154 | 0.0950 | 0.0540 | 0.0611 | 0.1377 |
| **Mot** | *Mean-centered* | -0.0276 | 0.0298 | 0.0022 | -0.0044 | 0.0429 |
| | *Non-centered* | -0.0105 | 0.0593 | 0.0020 | -0.0044 | 0.0480 |
| **Rel** | *Mean-centered* | -0.0164 | 0.0758 | -0.0227 | 0.0496 | 0.0547 |
| | *Non-centered* | 0.0721 | 0.0358 | -0.0228 | 0.0496 | 0.0658 |
| **Soc** | *Mean-centered* | 0.0085 | 0.0728 | 0.0071 | 0.02 | 0.0619 |
| | *Non-centered* | 0.0403 | 0.0034 | 0.0059 | 0.0199 | 0.0431 |
| **WM** | *Mean-centered* | -0.0111 | 0.0938 | 0.0236 | 0.1323 | 0.1131 |
| | *Non-centered* | 0.0626 | 0.0573 | 0.0248 | 0.1322 | 0.0992 |

**Supplementary Table 11.** Mean prediction performance of rCPM (5 iterations, *P* threshold = 0.1) performed on PPI betas generated after mean centering all predictors ("Mean-centered") and without centering the task regressors (i.e., condition on = 1, condition off = 0; "Non-centered"). Results presented as mean $q^2$.

| | Intercept | ciFC | cdFC | Activation | Combined |
|---|---|---|---|---|---|
| **Emotion** | -0.0082 | 0.0246 | -0.0428 | -0.0055 | -0.0041 |
| **Gambling** | 0.0355 | 0.0303 | -0.0358 | -0.007 | 0.0335 |
| **Language** | 0.0757 | 0.0689 | 0.0449 | 0.0474 | 0.1136 |
| **Motor** | -0.0272 | 0.0169 | 0.0083 | -0.0044 | 0.0420 |
| **Relational** | 0.0181 | 0.0750 | -0.0198 | 0.0547 | 0.0558 |
| **Social** | 0.0271 | 0.0349 | 0.0125 | 0.0238 | 0.0562 |
| **WM** | 0.0416 | 0.077 | 0.0049 | 0.0999 | 0.0944 |

**Supplementary Table 12.** Mean prediction performance of rCPM (5 iterations, *P* threshold = 0.1) performed only on the subset of participants who did not experience HCP QC issue C or recon version r177 ($n$ = 531). While prediction performance suffered overall due to the decreased sample size, relative performance patterns are comparable to those in main results (see main text).
